# High-resolution view of the type III secretion export apparatus *in situ* reveals membrane remodeling and a secretion pathway

**DOI:** 10.1101/709592

**Authors:** Carmen Butan, Maria Lara-Tejero, Wenwei Li, Jun Liu, Jorge E. Galán

## Abstract

Type III protein secretion systems are essential virulence factors for many important pathogenic bacteria. The entire protein secretion machine is composed of several substructures that organize into a holostructure or injectisome. The core component of the injectisome is the needle complex, which houses the export apparatus that serves as a gate for the passage of the secreted proteins through the bacterial inner membrane. Here we describe a high-resolution structure of the export apparatus of the *Salmonella* type III secretion system in association with the needle complex and the underlying bacterial membrane, both in isolation and *in situ*. We show the precise location of the core export apparatus components within the injectisome and bacterial envelope and demonstrate that their deployment results in major membrane remodeling and thinning, which may be central for the protein translocation process. We also show that InvA, a critical export apparatus component, forms a multi-ring cytoplasmic conduit that provides a pathway for the type III secretion substrates to reach the entrance of the export gate. Combined with structure-guided mutagenesis, our studies provide major insight into potential mechanisms of protein translocation and injectisome assembly.

## Introduction

Type III protein secretion systems (T3SSs) have the remarkable capacity to deliver multiple bacterially encoded effector proteins into target eukaryotic cells [1–3]. They are central for the virulence of many important bacterial pathogens and therefore have been long considered prime targets for the development of novel anti-infective strategies [4–6]. T3SSs are evolutionary related to flagella and, consequently, some of its components share amino acid sequence or structural similarities with components of the flagellar apparatus [7].

The central element of the protein secretion machine is the injectisome, a multi-protein structure composed of the needle complex and the cytoplasmic sorting platform (Fig. 1A) [8, 9]. The needle complex is made up of a base structure, embedded in the bacterial envelope, and a filamentous extension or needle that projects several nanometers from the bacterial surface. The base is composed of a double-ring structure, inner ring 1 (IR1) and inner ring 2 (IR2), which are made up of PrgH and PrgK in the *Salmonella* pathogenicity island 1 (SPI-1) T3SS, and the outer rings (OR1 and OR2) and neck, which are made up of InvG [10, 11]. Buried within the needle complex lies the export apparatus, a complex of several membrane proteins that facilitate the passage of type III secreted substrates through the bacterial inner membrane [12]. The entire needle complex is traversed by a narrow (∼3 nm) channel, which serves as a conduit for the proteins transiting the type III secretion pathway. On the cytoplasmic side of the needle complex lies the sorting platform, a multiprotein complex that engages, sorts, and initiates substrates into the secretion pathway [9].

**Fig. 1.**
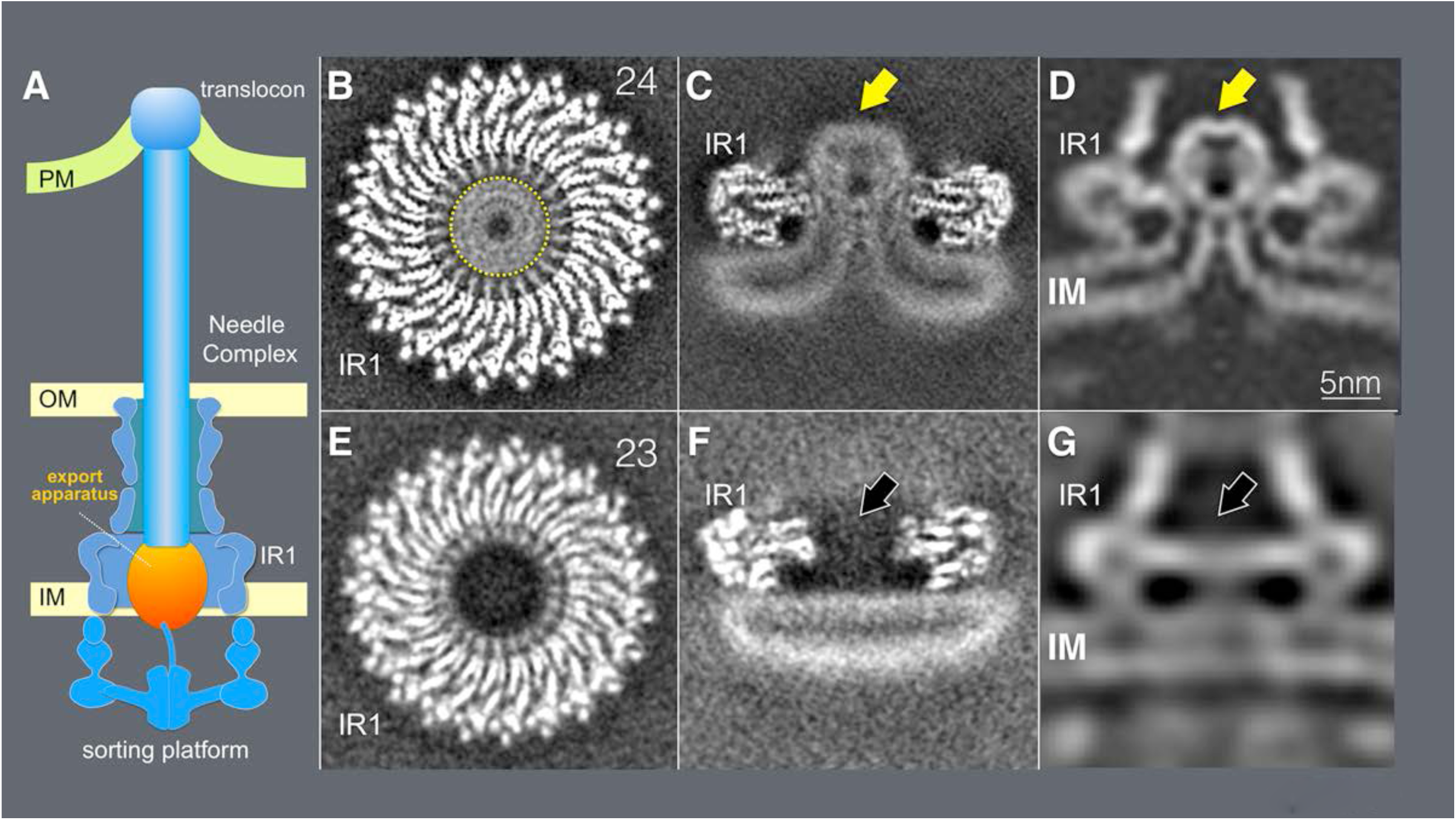
Cryo-EM and cryo-ET analyses of the core components the T3SS export apparatus. **(A)** A schematic representation of the T3SS injectisome. (**B** and **C**) Horizontal (**B**) and vertical (**C**) sections of a 3D reconstruction of a cryo-EM structure of the inner rings of the needle complex containing the export apparatus (denoted with a yellow arrow). (**D**) A vertical section from an *in situ* cryo-ET structure of the injectisome from a *S*. Typhimurium *ΔinvA* mutant strain. The export apparatus is denoted with a yellow arrow. (**E** and **F**) Horizontal (**E**) and vertical (**F**) sections of a 3D reconstruction of a cryo-EM structure of the inner rings of the needle complex obtained from a S. Typhimurium mutant strain *(ΔspaP ΔspaQ ΔspaR ΔspaS*) lacking the core components of the export apparatus. (**G**) A vertical section from in situ cryo-ET structure of the injectisome from a *ΔspaP ΔspaQ ΔspaR ΔspaS S*. Typhimurium strain. The position where the export apparatus would be located is denoted with a black arrow. IR1: inner ring 1; IM: inner membrane.

The complexity of the injectisome dictates that its assembly must occur in a highly coordinated, step-wise manner that is initiated by the formation of a complex of the core membrane protein components of the export apparatus, which are thought to nucleate the assembly of the lower rings [12, 13]. Following the recruitment of the independently assembled outer rings, the assembly of the injectisome is completed by the recruitment of the cytoplasmic sorting platform and the polymerization of the needle filament [14–16].

Previous studies have provided structural information on the different substructures that make up the injectisome [10, 11, 17–25]. However, there are still significant knowledge gaps, and importantly, it remains unclear how some of the structural information generated with isolated sub-complexes may ultimately relate to the injectisome structure *in situ*. This is particularly the case for the export apparatus, which is composed of five predicted membrane proteins (SpaP, SpaQ, SpaR, SpaS, and InvA in the *S.* Typhimurium SPI-1 T3SS also known as SctR, SctS, SctT, SctU, and SctV in the proposed unifying nomenclature) that are highly conserved in both, the virulence-associated type III secretion and flagellar assembly systems [12, 26]. A recent single particle cryo electron microscopy (cryo-EM) structure of a FliP/FliQ/FliR complex of the flagellar export apparatus (homologs of SpaP/SpaQ/SpaR) surprisingly showed that, in isolation, these proteins do not adopt typical integral membrane protein topologies but rather, they organize in a helical assembly that was proposed to be located largely within the periplasmic portion of the secretion machine [27]. Therefore it remains unclear how this complex facilitates the passage of type III secreted proteins through the inner membrane. Even less structural information is available about InvA [28], which is thought to play a central role in energizing the secretion process [29]. Although it has been proposed that the cytoplasmic domain of this protein family forms a nonameric ring interfacing with components of the sorting platform and export apparatus [25, 30], the localization of its critical transmembrane domain region within the secretion machine has remained elusive.

In this study, we have utilized single-particle cryo-EM and cryo-electron tomography (cryo-ET) to obtain a high resolution view of the export apparatus of the *S.* Typhimurium T3SS encoded within its pathogenicity island 1 (SPI-1), both *in situ* and in the context of needle complexes obtained with an isolation protocol optimized to maintain its native structural organization. These approaches have allowed us to determine the precise topological organization of the export apparatus relative to other components of the type III secretion injectisome, to characterize its interface with the base substructure, and to reveal major membrane remodeling concomitant with the assembly of the export apparatus. Furthermore, we have defined the structural organization of InvA, which outlines a multi-ring conduit for the type III secretion substrates to reach the entrance of the export apparatus gate. These studies provide major insight into the structure and assembly of the type III secretion injectisomes and suggest a pathway for the type III secreted substrates to cross the bacterial membrane.

## Results

### Cryo-EM and cryo-ET analysis of the core components the T3SS export apparatus and associated structures reveals major remodeling of the bacterial inner membrane

Previous structural studies of isolated T3SS injectisomes have relied on isolation procedures that result in the loss of some of its components or the disruption of its interaction with surrounding structures [10, 11, 24]. To address this limitation, we developed a needle complex isolation protocol under mild conditions that results in minimal losses of export apparatus components. Extraction of the needle complex from the bacterial envelope under these mild conditions was facilitated by the removal of the outer rings and neck components, which anchor this structure to the bacterial outer membrane (see Materials and Methods). LC-MS/MS analysis of the isolated structures detected the presence of the lower ring components PrgH and PrgK, the inner rod and filament proteins PrgJ and PrgI, as well as the export apparatus components SpaP, SpaQ, and SpaS (S1 Table). The apparent absence of the core component of the export apparatus SpaR was likely due to its lower stoichiometry in the complex [27, 31] and difficulties in its detection by LC-MS/MS under the conditions used in our analysis. Although InvA was also present in the sample, only few peptides were detected despite its large size and high stoichiometry. This indicates that most likely, InvA was absent from most of the particles, which is consistent with previous observations suggesting its loose association to the core components of the export apparatus [12]. Cryo-EM grids with the isolated structures (S1 Fig.) were examined by single particle analysis as described in Materials and Methods. We found that the most abundant population of substructures consisted of inner rings with the associated export apparatus but lacking the needle filaments (Fig. 1 and S1 Fig.). However, inner rings with associated needle structures were also observed in a proportion (∼5%) of the total number of particles compiled from the electron micrographs (S1 Fig.). Reference-free two-dimensional (2D) class average images of the inner rings with the associated export apparatus revealed elements of secondary structure in the IRI (S1 Fig.). However, the IR2 was poorly defined in all 2D class averages (Fig. 1 and S1 Fig.), which is expected for this substructure that undergoes a significant repositioning upon its isolation leading to varying locations relative to the rest of the needle complex structure [25], thus hampering its visualization at high resolution. After subtraction of the less-resolved IR2 signal and imposing 24-fold symmetry the resolution of the IR1 could be further improved (S1 Fig.). Residues 171-364 of PrgH and 20-203 of PrgK could therefore be docked with an excellent fit into the densities of this ring (Fig. 2).

**Fig. 2.**
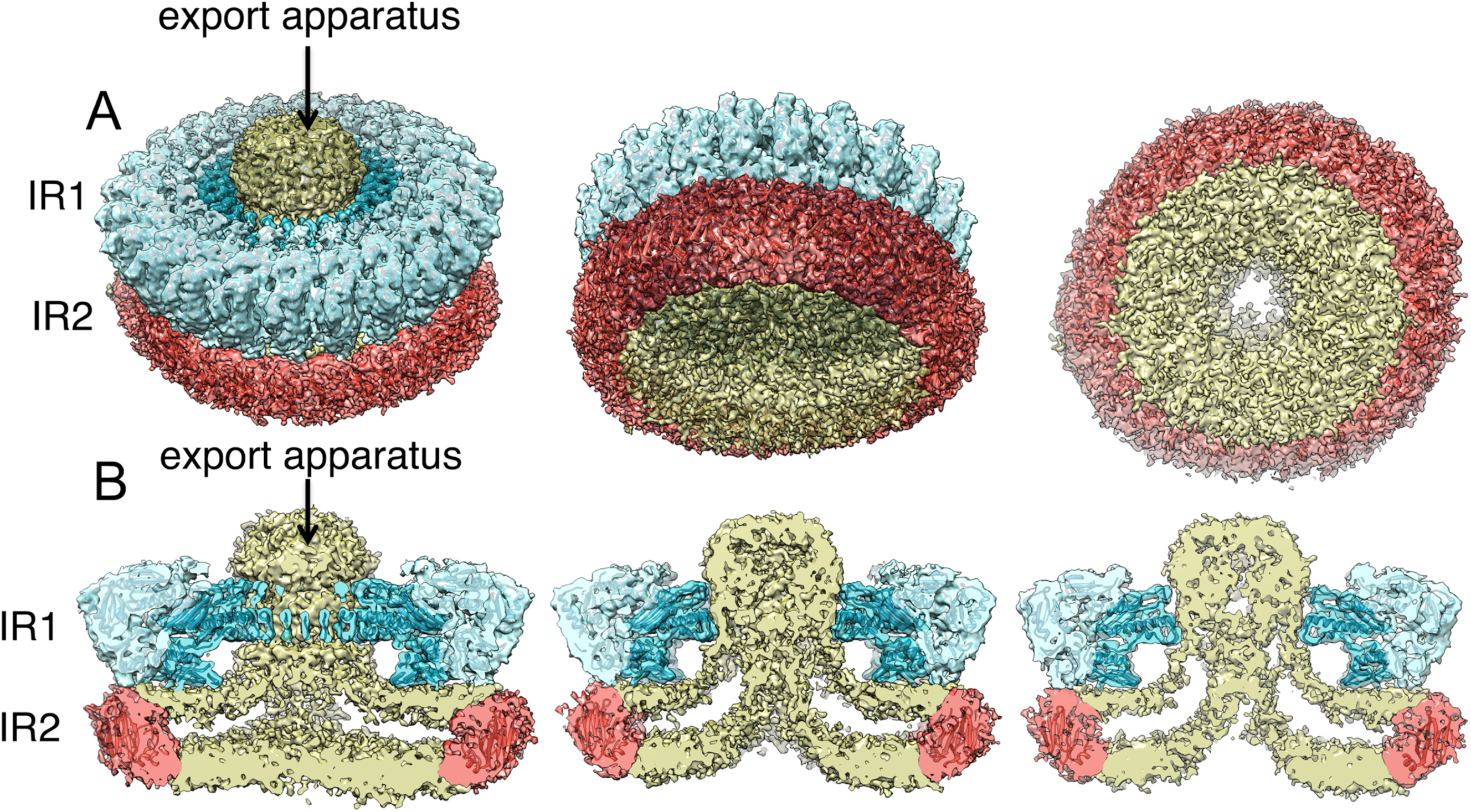
3D reconstructions of the needle complex IR1 and IR2 containing the core components of the export apparatus and associated membranes. (**A** and **B**) 3D density maps of the inner membrane rings containing the export apparatus with the crystal structures of PrgH (residues 171-364, light blue) and PrgK (residues 20-203, dark blue) docked into the density corresponding to the IR1. Although the resolution is not sufficient for confident assignment, a modeled structure of residues 11 to 120 of PrgH (red) docked into the density corresponding to the IR2 is shown to mark the expected position of this structural element. Tilted (**A**) and cut away (**B**) views are shown. Densities corresponding to the export apparatus and associated membranes are shown in yellow.

Further analysis of the “side views” in the 2D class averages of structures lacking the needle filament revealed a cloud of density spanning the space between IR1 and IR2 and extending above IR1 (Fig. 1C, 2A and 2B, and S1, S2 and S3 Fig.). We assigned this cloud of density to the core protein components of the export apparatus since this density was absent in structures isolated from an isogenic *ΔspaP ΔspaQ ΔspaR ΔspaS S*. Typhimurium mutant strain (Fig. 1E, 1F and S4 Fig.). The 2D class averages of tilted views of the rings and the 3D reconstruction of structures isolated from this *S*. Typhimurium mutant strain showed an empty space at the center of the double ring structure (Fig. 1E, 1F and S4 Fig.).

**Fig. 3.**
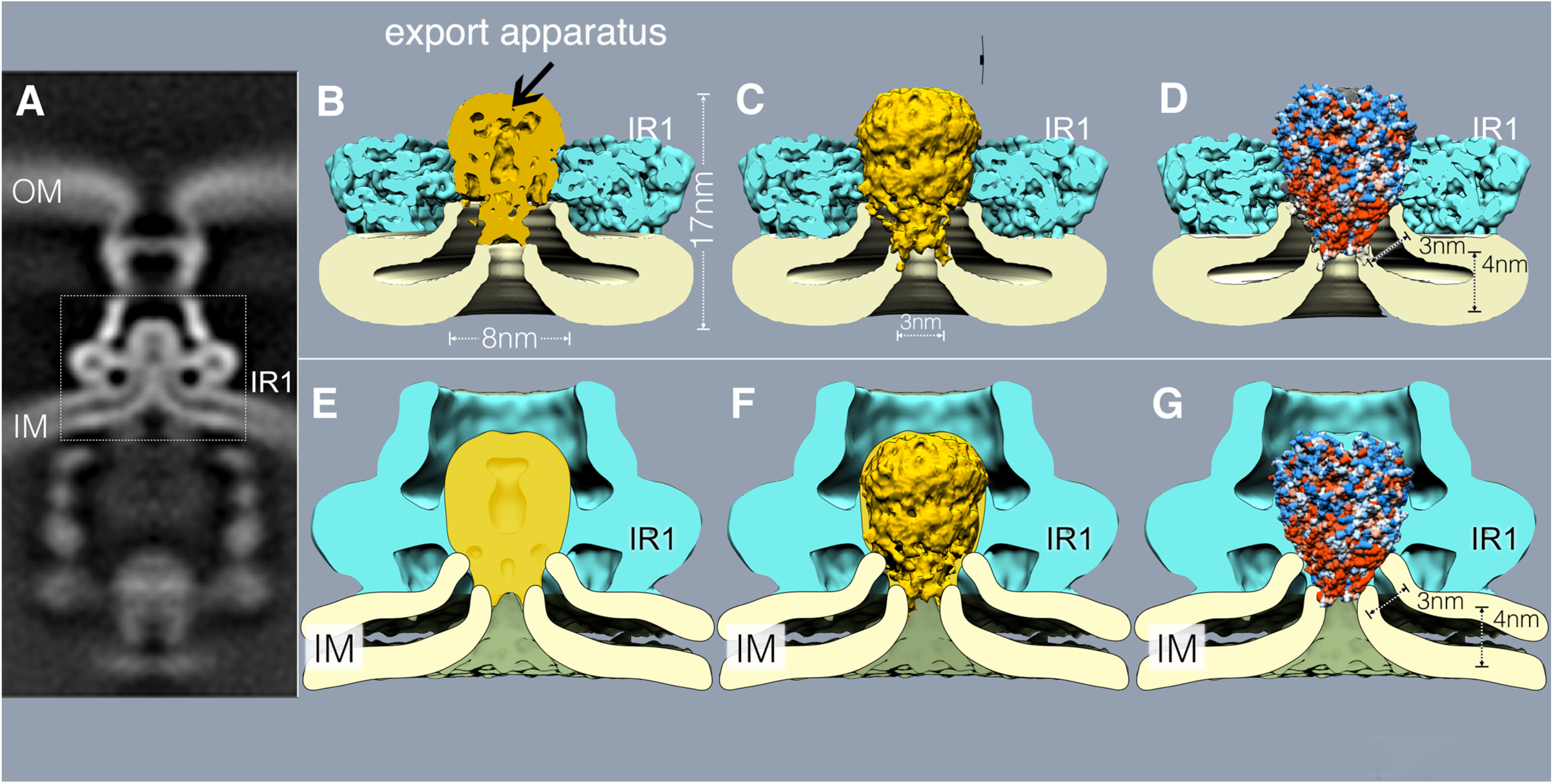
Assembly of the export apparatus results in pronounced membrane remodeling. (**A**) A vertical section from *in situ* cryo-ET structure of the injectisome from a *ΔinvA S*. Typhimurium strain (the dotted box denotes area of detail shown in **B** through **G**). (**B-G**) Vertical sections of a cryo-EM (**B**-**D**) and cryo-ET (**E**-**G**) structures of the inner rings of the needle complex T3SS complex overlaid with a section (**B** and **E**), surface rendering (**C** and **F**) or hydrophobicity surface representation (**D** and **G**) of the FliP/FliQ/FliR (homologs of SpaP/SpaQ/SpaR from *S*. Typhimurium) isolated complex (PDB 6F2E). Red and blue colors (**D** and **G**) on the surface representation of the FliP/FliQ/FliR isolated complex represent hydrophobic and charge residues, respectively.

**Fig. 4.**
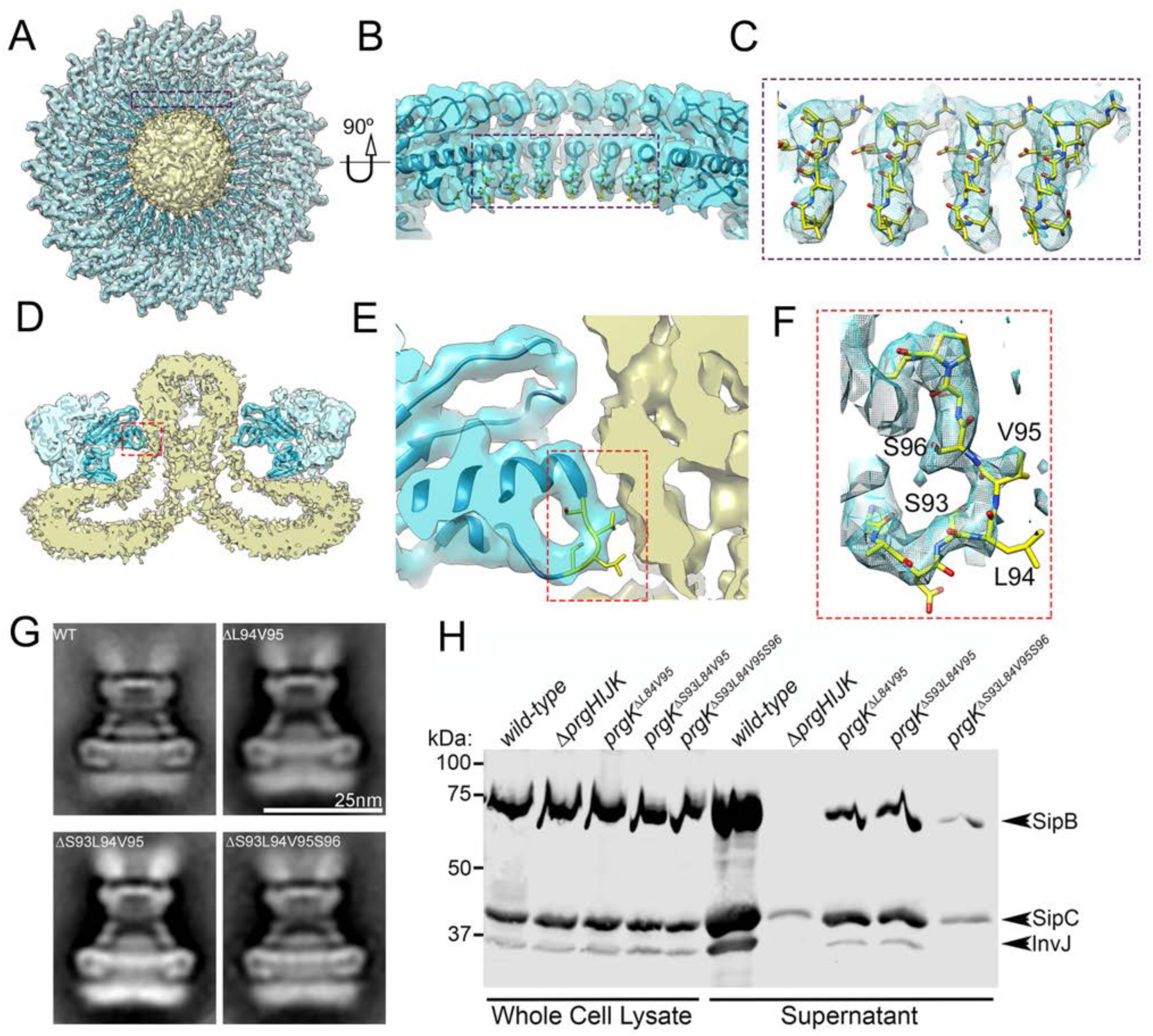
Interaction of the periplasmic PrgK ring with the export apparatus is required for type III secretion function but not for injectisome assembly. **(A - F)** Top view (**A**) and cross section (**D**) of the 3D reconstruction of the IR1 and IR2 rings of the needle complex containing the core components of the export apparatus. The interface between PrgK and the export apparatus is highlighted (in blue and red rectangles). The specific densities corresponding to PrgK residues 93-96 targeted for mutagenesis and functional analysis are shown in more detailed in the zoom-in views depicted in panels **B** through **F**. (**G**) Representative class averages of needle complexes isolated from S. Typhimurium expressing wild type PrgK or the indicated deletion mutants. (**H**) Type III protein secretion analysis of S. Typhimurium strains expressing the PrgK mutants with the indicated deletions in the region that interacts with the core components of the export apparatus. Whole cell lysates and cultured supernatants of the indicated strains were analyzed by Western immunoblot for the presence of SipB, SipC, and InvJ, which are substrates of the S. Typhimurium type III secretion system.

Comparison of this inner ring substructure with previously reported needle complex structures showed a notable difference, characterized by the presence of a large double-layered density surrounded by the IR2 that occupies the entire cytoplasmic-side of the double ring structure (Fig. 1 and 2, and S5 Fig.). This density, which cannot be accounted for by the cytoplasmic domain of PrgH (Fig. 2), forms a funnel-like structure with its center leading to the entrance of the export apparatus (Fig. 1 and 2, and S5 Fig.). The overall thickness (∼4 nm) and double layer appearance of this density indicates that this structure corresponds to the bacterial inner membrane, which under our mild isolation conditions, has been retained in the isolated structures. Consistent with this hypothesis, the 3D reconstruction of structures isolated from the strain lacking the export apparatus showed that in this mutant, the double layer density no longer organizes in a funnel shape but rather, appears flat completely sealing the space surrounded by the IR2 (Fig. 1F, and S4 Fig.). These results indicate that the deployment of the export apparatus results in a rather significant remodeling of the surrounding bacterial membrane.

**Fig. 5.**
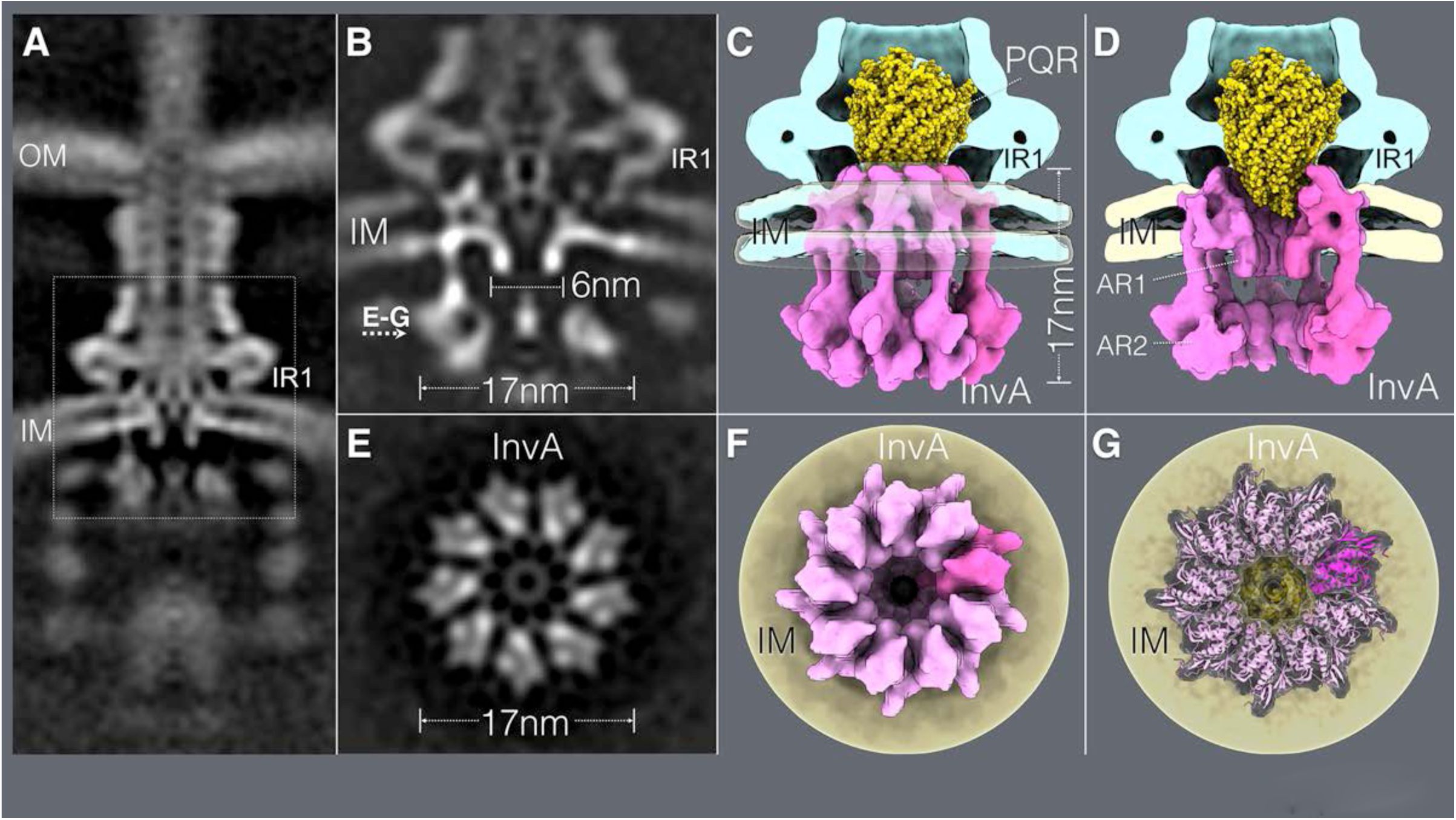
*In situ* structure of InvA, an essential component of the type III secretion export apparatus. **(A)** A vertical section from the *in situ* cryo-ET structure of the injectisome from wild-type *S*. Typhimurium (the dotted box denotes area of detail shown in **B** through **G**). (**B - G**) Vertical (**B**-**D**) and horizontal (through the plane indicated in panel B) (**E**-**G**) sections highlighting the structure of InvA. Side (**C**) and cut-away (D) views of the InvA structure (in pink) are shown and the location of the small (AR1) and large (AR2) rings are denoted. The overlaid of the structure of the FliP/FliQ/FliR isolated complex is shown in yellow. (**F** and **G**) Bottom views of the InvA nonameric ring, with the overlaid atomic structure of the cytoplasmic domain of an InvA homolog (PDB-4A5P) (**G**).

To ascertain whether the membrane remodeling observed in isolated structures is also present in *in situ* structures embedded within the bacterial envelope, we examined by cryo-ET, injectisomes in bacterial minicells obtained from an *S*. Typhimurium strain lacking *invA* (which was absent in our samples thus better resembling the isolated complex). Subtomogram average from 5,929 injectisome reconstructions shows a rather pronounced rearrangement of the inner membrane in the vicinity of the density corresponding to the export apparatus (Fig. 1D). Like in the isolated complex, in the *in situ* structure the membrane is seen adopting a funnel-like shape with its narrow end in close proximity to the density that corresponds to the export apparatus. The remodeled membrane was not observed in *in situ* structures obtained from a mutant lacking all the components of the export apparatus (subtomogram average from 2,368 injectisomes reconstructions) (Fig. 1G), indicating that membrane remodeling is a consequence of the deployment of the core components of the export apparatus. Importantly, the two isolated cryo-EM injectisome structures and associated membranes could be overlaid onto the cryo-ET structures with an excellent fit (Fig. S6), indicating that the isolation procedure did not disrupt the topology of the export apparatus relative to the surrounding membranes and injectisome substructures.

Taken together, these results indicate that the mild isolation protocol resulted in needle complex substructures that have retained the export apparatus and associated membranes, thus maintaining the *in situ* topology. More importantly, these observations indicate that the bacterial inner membrane undergoes a significant remodeling upon deployment of the core components of the export apparatus.

### Needle complex inner rings assembled in the absence of the export apparatus exhibit a different symmetry

The 2D class averages and the 3D reconstruction of structures obtained from the *S.* Typhimurium mutant strain lacking the export apparatus showed that the vast majority (∼80%) of the inner membrane rings exhibited 23-fold symmetry (Fig. 1E Fig. S4 Fig.). This observation is significant since it is in marked contrast to the inner membrane rings assembled in the presence of the export apparatus, which invariably exhibits 24-fold symmetry (Fig. 1B and S1 and S2 Fig.). We have previously shown that the efficiency of assembly of the needle complex is markedly reduced in *S*. Typhimurium mutants lacking the export apparatus, which in combination with additional biochemical data led us to propose that the export apparatus templates the assembly of the inner rings [12]. The observation that inner rings assembled in the absence of the export apparatus exhibit symmetry never observed in the rings assembled in its presence is consistent with this model and strongly supports the notion that during assembly of a functional injectisome the export apparatus must be deployed prior to the assembly of the inner rings.

### Overall architecture of the core components of the export apparatus in association with the inner rings and the bacterial membrane

Overall, the density associated with the export apparatus in the context of needle complex rings is ∼11 nm long from top to bottom and ∼8 nm at its widest (S7 Fig.). The cryo-EM structure of an isolated sub-complex of the flagellar-associated export apparatus made of FliP, FliQ and FliR (homologs of SpaP, SpaQ, and SpaR, respectively) [27] revealed that none of its subunits adopt a typical membrane protein topology, but rather they participate in the formation of a helical structure that was postulated to be largely located in the periplasm [27]. This observation raised the question how this complex may insert in the membrane, an essential requirement to mediate the passage of secreted proteins through the membrane barrier. Analysis of the surface hydrophobicity of the FliP/FliQ/FliR complex revealed only a small hydrophobic strip at the base of the structure, which was hypothesized to be buried in the membrane [27] (S8 Fig.). The length of this segment (∼30 Å) would predict that such hydrophobic region would not expand a typical membrane thus raising intriguing questions about the mechanism by which this complex may mediate protein translocation across the membrane. Although attempts were made to place the isolated structure onto a low-resolution cryo-ET map of the export apparatus [27], it has remained unclear what the precise location of the core component of the export apparatus relative to the bacterial inner membrane is. We therefore docked the a cryo-EM structure of the isolated FliP/FliQ/FliR complex of the flagellar-associated export apparatus into our structures, which allowed us to precisely place this export apparatus sub-complex in its appropriate topological context. Overall, there is good agreement between the FliP/FliQ/FliR isolated complex structure and the cryo-EM and cryo-ET structures of the *S*. Typhimurium export apparatus embedded within the needle complex, particularly in the densities that correspond to the periplasmic domain of the export apparatus (Fig. 3B-3G and Fig. S8). Close examination of the overlay of the structures reveals that the hydrophobic vertex of the FliP/FliQ/FliR complex is predicted to be located at the expected plane of the membrane (Fig. 3B-3G and S8 Fig.). Importantly, both our cryo-EM and cryo-ET structures suggest that the substantial remodeling of the inner membrane associated with the deployment of the export apparatus results in a significant membrane thinning in the immediate vicinity of the export apparatus (Fig. 3B-3G and S8 Fig.). The thinned membrane allows the hydrophobic vertex of the export apparatus core complex to expand the membrane in its entire width (Fig. 3B-3G, S8 Fig., and S1 Video). This architecture places the cavity observed at the vertex of the cryo-EM structure of the FliP/FliQ/FliR complex in direct communication with the cytoplasm of the bacterial cell (Fig. 3B-3G, S8 Fig., and S1 Video), an arrangement that may facilitate the passage of the substrates through the membrane. These observations have fundamental implication for the understanding of the protein translocation mechanism across the bacterial inner membrane during type III secretion.

### Interactions of a domain of the periplasmic PrgK ring with the core components of the export apparatus are required for type III secretion

Close examination of the interface between the needle complex IRI and the export apparatus revealed a segment of PrgK that makes the most intimate contact with the periplasmic region of the export apparatus (Fig. 4A-4F). This region consists of a 6 amino-acid loop (residues 92 to 98), which is anchored by two helixes (Fig. 4F). To examine the functional significance of this intimate association between IR1 and the export apparatus we introduced discrete mutations in this loop region of PrgK and examined the structure and function of the needle complex. We found that structures isolated from *S*. Typhimurium strains expressing PrgK^Δ94-95^, PrgK^Δ93-95^, or PrgK^Δ93-96^ were indistinguishable from wild type except for the reduced number of needle filaments associated to these structures, which was particularly apparent in the structures obtained from the strain expressing the PrgK^Δ93-96^ mutant (Fig. 4G and S9 and S10 Fig.). The export apparatus was readily detectable in the needle complexes isolated from all mutants, which at this level of resolution appeared indistinguishable from wild type (Fig. 4G and S9 Fig.). Despite the apparently normal appearance of the injectisome, all the mutants showed a marked decrease in type III secretion function as measured by the ability of these *S*. Typhimurium *prgK* mutant strains to secret various substrates of the SPI-1 T3SS (Fig. 4H). The mutant phenotype was particularly apparent for the *prgK^ΔS93-S96^* mutant strain, which showed almost complete loss of function (Fig. 4H). These observations indicate that this intimate interaction between PrgK and the export apparatus is central for type III secretion but not for the assembly of the needle complex and the deployment of the export apparatus. These observations suggest an active role for the PrgK ring substructure in T3SS function beyond the scaffolding of the export apparatus in the bacterial membrane.

### *In situ* structure of InvA, a highly conserved component of the export apparatus

InvA (FlhA in the flagellar system and SctV in the proposed unifying nomenclature) is one of the most highly conserved components of the export apparatus [28, 32]. However, its role in the protein secretion process is poorly understood. Secondary structure predictions and molecular modeling indicate that the amino terminal half of this protein family contains several transmembrane segments and its large carboxy-terminal domain is located entirely within the bacterial cytoplasm [32, 33]. It has been proposed that this protein family, alone or in association with other components of the export apparatus, may function as a proton or Na+ channel that contributes to energizing the secretion process [34, 35]. However, how this protein may contribute to this activity is still uncertain and it is unclear whether it forms a physical complex with other components of the export apparatus. Of critical importance for the understanding of the function of this crucial component of the T3SS is to identify the location of the predicted transmembrane region within the context of the injectisome structure. Efforts to isolate InvA (or any of its homologs) in association with the injectisome have failed. The crystal structure of a region of the carboxyterminal cytoplasmic domain of a member of this protein family suggests that it can forms a nonameric ring [30], which previous cryo-ET imaging has located in the bacterial cytoplasm, in close apposition to components of the sorting platform [25]. However, there is no structural information on the critical amino terminal half of this protein, which is responsible for positioning it within the structure of the injectisomes. We therefore applied a high-throughput cryo-ET pipeline to image minicells obtained from a *S.* Typhimurium strain expressing a wild type SPI-1 T3SS and applied sub-tomogram averaging to image InvA. We found that InvA forms a sea-horse-like structure, with the head embedded in the membrane and the body and tail located entirely within the cytoplasm (Fig. 5A-5D and S1 Video). Consistent with the crystal structure [30], the cytoplasmic domain of InvA forms a nonameric ring 17 nm in diameter and 5.5 nm in height (Fig. 5). This ring is enclosed within the sorting platform immediately above a density we have previously assigned to the ATPase InvC [25]. Importantly, immediately above this ring and closer to the membrane but still within the bacterial cytoplasm, we detected an additional ring 6 nm in diameter and 8 nm in height (Fig. 5 S2 Video). This smaller ring aligns very well with the larger ring and marks the entrance to a broad conduit that narrows as it gets closer to the export apparatus gate. This conduit is bounded by densities that we postulate correspond to the transmembrane domains of InvA, whose arrangement tracks the contour of the drastically remodeled bacterial membrane. Consistent with this notion, subtomogram averages of injectisome from an *S.* Typhimurium strain expressing an InvA mutant lacking its last 329 amino acids, which removes its entire cytoplasmic domain, lack the larger cytoplasmic ring but retained the smaller ring and membrane proximal structures (S11 Fig.). None of these structures were present in a strain lacking InvA (Fig. 1D) although they were readily detected in a strain lacking the export apparatus component SpaS (S11 and S12 Fig.). All combined, the two rings and membrane-associated densities build a contiguous substructure that we hypothesize initiates the substrates of the type III secretion machine into the secretion pathway (S2 Video). Of note, sagittal sections through the injectisome cutting through the PrgH cytoplasmic IR2 domain showed an intimate association between this structure of the needle complex and a surface region (dubbed SD2 in previous studies [30]) within the larger InvA cytoplasmic ring (Fig. 6). Discrete mutations within this region of a homolog of InvA affected type III secretion function without affecting the assembly of the InvA ring [30]. Consequently, our observations may explain this phenotype suggesting a potential mechanism by which conformational changes in the needle complex could modulate the activity of InvA, which plays a central role in the initiation of substrates into the secretion pathway.

**Fig. 6.**
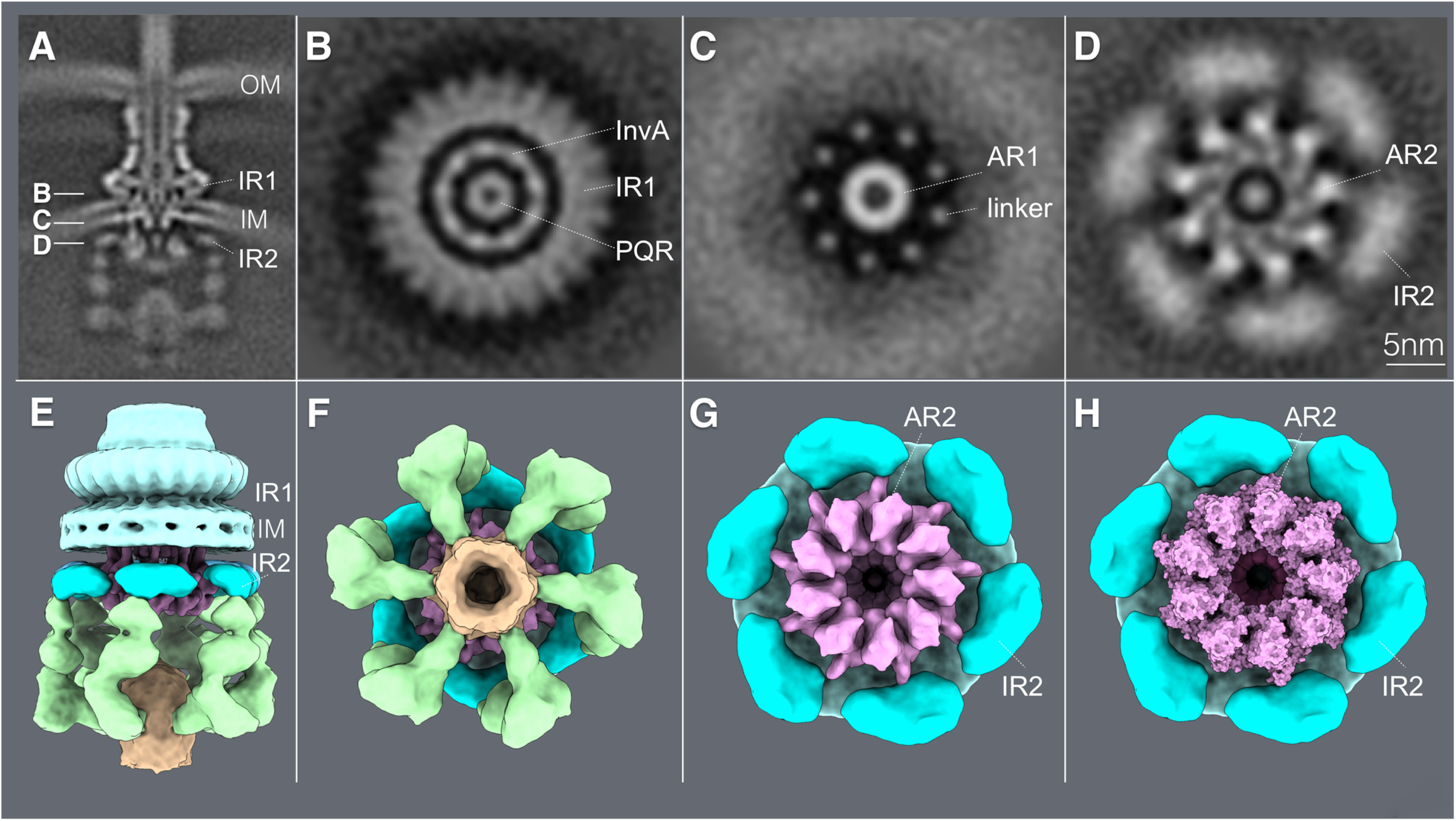
Close association of a cytoplasmic ring of InvA with the sorting platform and cytoplasmic domain of the needle complex component PrgH. (**A**) A vertical section from the *in situ* cryo-ET structure of the injectisome from wild-type *S*. Typhimurium indicating the section planes of the structures shown in (**B**-**D**). (**B**-**D**) Horizontal sections through the planes indicated in (**B**) depicting the InvA nonameric ring within the inner ring 1 (IR1) of the needle complex structure (**B**), the linker region that joins the membrane and cytoplasmic domains of InvA (**C**), and its tight association with the cytoplasmic domain of the inner ring 2 (IR2) of the needle complex (**D**). The position of the core component of the export apparatus (labeled PQR) is also shown. (**E and F**) Side (**E**) and bottom (**F**) views of the surface rendering of the cytoplasmic elements of the T3SS injectisome. (G) Bottom view of the T3SS injectisome after removal of the sorting platform elements to highlight the close association between the larger cytoplasmic ring of InvA (AR2) with the cytoplasmic ring of PrgH (IR2). (**H**) Bottom view of the InvA nonameric ring (AR2) with the overlaid atomic structure of the cytoplasmic domain of an InvA holog (PDB-4A5P).

## Discussion

The export apparatus is a central core component of T3SS that is thought to mediate the passage of type III secreted substrates through the bacterial membrane, a critical step during the protein translocation process. All the core components of the export apparatus (SpaP, SpaQ, SpaR, SpaS, and InvA in the *S*. Typhimurium SPI-1 T3SS) are predicted to be integral membrane proteins [33]. However, a recent cryo-EM structure of an isolated sub-complex of the flagellar-associated export apparatus made up of FliP, FliQ, and FliR (homologs of SpaP, SpaQ and SpaR) revealed that none of these proteins adopt a typical membrane protein topology and instead, they assemble into a helical structure that through fitting into a low resolution *in situ* structure of the injectisomes was proposed to be largely located within the bacterial periplasm [27]. The absence of typical transmembrane regions in this sub-complex is intriguing in light of their proposed role in mediating the passage of type III secreted proteins through the bacterial membrane, and raised the question whether other components of the export apparatus, could fulfill this role. To address these issues we have utilized single particle cryo-EM and cryo-ET to determine the precise architecture of the export apparatus *in situ* and, in particular, to determine the topological organization of its components relative to the bacterial membrane.

To this aim, we developed an isolation protocol that allowed us to obtain needle complex substructures containing the core components of the export apparatus in association with the bacterial membrane. Analysis of this structure by single-particle cryo-EM allowed us to determine the precise topological relationship between the core components of the export apparatus, the needle complex, and the associated bacterial membrane. Notably, we found that the bacterial membrane undergoes a profound remodeling in the immediate vicinity of the export apparatus. This remodeling, which is also observed in the cryo-ET *in situ* structure, is characterized by a significant outward “pinching” of the membrane, which organizes in a “funnel like” fashion with its narrower end in immediate contact with the core components of the export apparatus. Notably, this reorganization was not observed in injectisome sub-complexes isolated from a Δ*spaPΔspaQΔspaR*Δ*spaS S. Typhimurium* mutant strain, indicating that membrane remodeling is strictly dependent on the deployment of the core components of export apparatus. The membrane remodeling was also accompanied by a significant thinning of the bacterial membrane, particularly in the immediate vicinity of its interface with the export apparatus. The thinning of the membrane may allow the relatively short (∼30 Å) hydrophobic vertex of the core complex of the export apparatus to expand the entire membrane thus facilitating the movement of the type III secreted proteins through the bacterial inner membrane. Furthermore, this topological organization would allow a central cavity present within this vertex to be in direct communication with the bacterial cytoplasm thus facilitating substrate translocation across the membrane. These observations have important implications for the understanding of the mechanisms involved in the protein translocation function of the export apparatus as it implies that this complex would have all the necessary structural elements to form the channel that mediate the passage of the secreted substrates through the bacterial inner membrane. It is noteworthy that the thinning of the membrane and a similar topological organization have also been observed in the context of other protein conducting channels such as the Hrd1, twin arginine translocator (TAT), and YidC protein translocation machines [36–38], suggesting that this type of membrane remodeling may be a more general feature associated with some protein translocation machines.

We have identified in our structure a region of PrgK, an inner ring component of the needle complex, which makes intimate contact with the periplasmic region of the core complex of the export apparatus. Introduction of discrete mutations in this domain resulted in loss of function, even though these mutations did not detectably affect the assembly of the PrgK rings or the deployment of the export apparatus. It is therefore possible that PrgK may trigger conformational changes in the export apparatus that may be necessary to “prime” its secretion function. These observations point to an unexpected active role of the PrgK rings in the type III secretion process, presumably involving more than just serving as a scaffold for the export apparatus.

We have previously shown that in the absence of the export apparatus the efficiency of assembly of the needle complex is significantly diminished, which in combination with other observations led us to propose an injectisome assembly pathway that is initiated by the deployment of the core components of the export apparatus [12]. However, alternative models of assembly have also been proposed [16]. We have shown here that needle complexes assembled in the absence of the export apparatus exhibit a fold symmetry that is never observed in fully assembled needle complexes. Therefore, this observation demonstrates that the assembly of the core components of the export apparatus must be the first step in the injectisome assembly pathway.

Despite their central role in protein secretion, the mechanisms by which the InvA protein family contributes to type III secretion has remained elusive. Although it has been proposed to exert its function as a proton channel that energizes the secretion process [34], direct demonstration of its proton transport activity is lacking. Furthermore, it is unclear how a putative proton channel activity contributes to the secretion process. A major limitation for the understanding of InvA function has been the lack of information on its precise localization within the secretion apparatus. The crystal structure of the carboxy terminal domain of InvA forms a nonameric ring, and this structural feature has been proposed to correlate with a toroidal density observed within the cytoplasmic sorting platform of the injectisome [25, 30]. However, there has been no information on the localization of the amino terminal half of InvA, which contains multiple predicted transmembrane domains. Here using a high throughput cryo-ET pipeline we have precisely determined the topological organization of the entire InvA molecule. Our *in situ* structure indicates that, as predicted by the crystal structure [30], the carboxy terminal domain of InvA (amino acids 357 to 685) forms a nonameric ring (17 nm in diameter) located in the apical side of the sorting platform cavity. In addition, our *in situ* structure revealed that InvA forms a second cytoplasmic ring 6 nm in diameter that is located in close apposition to the bacterial membrane and in alignment with the larger ring. We hypothesize that this ring is formed by a large cytoplasmic loop (amino acids 131 to 196) predicted by secondary structure and molecular modeling within the amino terminal half of InvA [33]. Consistent with this hypothesis, removal of the large cytoplasmic domain of InvA did not affect the localization and organization of its membrane proximal ring. Our structure also detected densities that most likely represent the membrane-embedded domain of InvA, which are located immediately adjacent (though apparently not in intimate contact) to the core components of the export apparatus, and within the remodeled funnel-shaped inner membrane. These observations have significant implications for the understanding of InvA function. First, the observed topological organization does not support a direct role for the transmembrane domains of InvA in the building of the putative protein translocation channel. This is significant since the lack of an obvious transmembrane regions in the SpaPQR complex that would be able to expand the entire membrane had raised the possibility that this complex may work in conjunction with InvA to form such a channel [27]. These observations coupled to the observed thinning of the membrane around the export apparatus strongly suggest that the SpaPQR complex (perhaps with SpaS) may be sufficient to form such channel as its entire vertex would be able to expand the thinned membrane. Rather, the topology of InvA raises the exciting possibility that through its series of cytoplasmic rings, InvA “guides” the substrates to the entrance of the export apparatus channel (Fig. 7 and S3 Video), a function that could be aided by its putative proton channel activities.

**Fig. 7.**
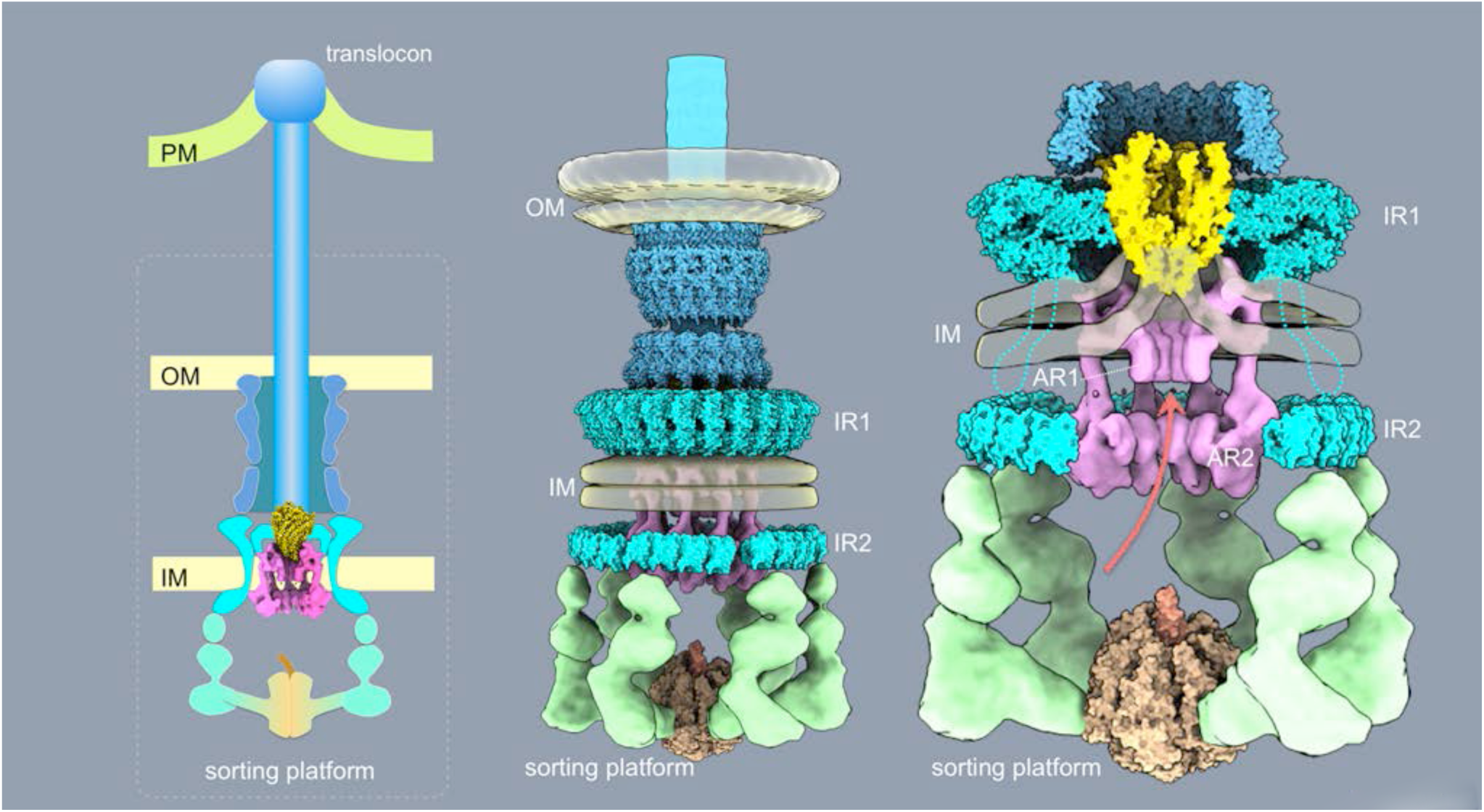
Schematic model of the export apparatus in the context of a fully assembled injectisome structure and the different bacterial envelope elements. PM: target host cell plasma membrane; OM: bacterial outer membrane; IM: bacterial inner membrane; IR1 and IR2: inner rings 1 and 2 of the T3SS needle complex; AR1 and AR2: membrane proximal (AR1) and distal (AR2) cytoplasmic rings of InvA.

A central feature of virulence-associated T3SS is that their protein-secretion activity is triggered only upon contact with eukaryotic cells [39, 40]. Consequently, it is expected that machines visualized in isolation or *in situ* after growth in culture medium would be in the “close” or inactive conformation. Since the activation of the machine is triggered by contact with target eukaryotic cells, the activating signal is likely to emerge from the tip structure of the needle filament and eventually transmitted to the export apparatus through the needle and inner rod substructures (i.e. “top down”) [41]. How this activating signal is relayed to the export apparatus is unclear. However, our observation that a discrete domain of the large cytoplasmic ring of InvA previously shown to be essential for type III secretion function [30] makes intimate contact with the IR2 cytoplasmic domain of the needle complex protein PrgH suggests a potential functional link between these two structures that may be important for transducing the activation signal. In this context, the close proximity of the export apparatus to both the transmembrane domains of InvA as well as a defined periplasmic domain of the needle complex protein PrgK (see above) may define potential sites of interactions that may serve as signal relay points to coordinate substrate engagement with the opening of the secretion channel. Furthermore, these signals may also lead to the activation/coordination of the putative proton motive force of InvA that may be required to advance the substrates through its multi-ring conduit that lead to the entrance of the export channel.

In summary, these studies have provided a close up view of the export apparatus in its *in situ* topological context and in association with other injectisomes components and the bacterial membrane. The topological relationship of these components to each other and to the bacterial membrane combined with the multi-ring organization of InvA suggest a potential pathway for the type III secreted substrates during their translocation through the bacterial envelope.

## Materials and Methods

### Bacterial strains and plasmids

All bacterial strains used in this study are derived from *Salmonella enterica* serovar Typhimurium strain SL1344 [42] and are listed in S2 Table. All strains were constructed by allelic exchange [43] and all plasmids were constructed by Gibson assembly [44]. The identity of all the strains was verified by nucleotide sequencing and relevant functional assays.

### Analysis of type III protein secretion function

The functionality of the SPI-1 T3SS in the *S*. Typhimurium strains was assessed by examining their ability to secrete type III secreted proteins to the culture supernatant as previously described. Briefly, overnight cultures were diluted 1/20 into LB containing 0.3M NaCl to induce the expression of SPI-1 T3SS [45]. Diluted cultures were grown at 37°C on a rotating wheel to an OD600 of ∼ 0.9 (4 to 5 hours), and the cells were pelleted by centrifugation at 6,000 rpm. Cells were resuspended in 1X SDS-running buffer and culture supernatants were filtered through a 0.45 µm syringe filter prior to the recovery of proteins by trichloroacetic acid (TCA) precipitation. Samples were subsequently analyzed by SDS-PAGE and Western blot with antibodies to the type III secreted proteins SipB, SipC and InvJ.

### Isolation of needle complexes substructures

The needle complex purification from the different S. Typhimurium strains (see Resource Table) was carried out as follows. An MBP-tagged PrgH allele, and when indicated a *ΔinvG* allele (to remove the outer rings and neck of the needle complex) were introduced into the different *S*. Typhimurium strains as indicated above. Two liters of 0.3M NaCl containing 100 µg/ml of ampicillin and 0.1% arabinose were inoculated with the different strains and grown for ∼10 hs under gentle (100 rpm) shaking. Cells were recovered by centrifugation at 6,000 rpm for 20 minutes, resuspended in 10 ml of lysis buffer [200mM Tris pH 7.5, 20% sucrose, 1mM EDTA, 0.25mg/ml of lysozyme and cOmplete™ EDTA-free protease inhibitor cocktail (Sigma 4693159001)] and incubated on ice for 1 hr. Cells were subsequently incubated for 5 min at 37°C and lysed by the addition of 0.5% N-Dodecyl- *ß-*D-maltoside (DDM) (Anatrace D310S). Cells were incubated at 37°C for additional 5 to 10 min while monitoring lysis, transferred to ice, and further incubated for 1 hr. Debris was removed by centrifugation at 14,000 rpm for 1 hour and the clarified lysate was transferred to a fresh tube. Two hundred microliters of amylose resin (NEB E8021) were added and the suspension was incubated O/N at 4°C under rocking conditions. Beads were then washed 4x with 10 ml of washing buffer (20 mM Tris pH 7.5, 100 mM NaCl, 1 mM EDTA) and finally resuspended in 50 µl of washing buffer containing 20 mM of maltose. After 1 hr incubation on ice with occasional tapping, beads were removed by centrifugation (3,000 rpm for 5 min) and the needle complex containing supernatant was transferred to a fresh tube for further analysis.

### Single-particle cryo-electron microscopy

Vitrified specimens were prepared by adding a 3.5 µl drop of the different samples to holey grids (TedPella, Quantifoil grids, Copper, 300 mesh, R1.2/1.3), which had been overlaid with an additional thin layer of carbon. Grids were blotted for 4-5 seconds, then flash-frozen in liquid ethane using a Vitrobot MarkIII or MarkIV instrument, with the chamber maintained at 6-7^0^C and 100% humidity. Grids were then imaged under two different conditions: (1) Grids overlaid with samples containing the inner membrane rings containing the export apparatus were imaged at the National Institute of Health (NIH) on a Titan Krios microscope (Thermo Fisher Scientific) operated at 300-kV cryo-electron microscope equipped with an energy filter (Gatan GIF Quantum), and a post-GIF Gatan K2 Summit direct electron detector. Before the imaging session the microscope was aligned for parallel illumination. Projection images were obtained with a K2 camera operated in super-resolution counting mode with a physical pixel size of 1.32 Å, and a super-resolution pixel size of 0.66 Å with the nominal defocus range set between −1.25 and −3.25 µm. During the image acquisition, the Gatan K2 Summit direct electron detector was operated in zero-energy-loss mode with a slit width of 20 eV. The dose rate was 4 e^-^/Å^2^/s at the specimen plane. The total exposure time was 12 s with intermediate frames recorded every 0.3 s, giving a total of 40 frames per image. The total accumulated dose was ∼48 e^-^/Å^2^; (2) Grids overlaid with samples containing the inner membrane rings without the export apparatus were imaged at the Yale West Campus on a Titan Krios microscope (Thermo Fisher Scientific) operated at 300-kV. Projection images were obtained using a Falcon III detector operated in counting mode with a physical pixel size of 1.07 Å and a nominal defocus range between −1.5 and −3.5 µm. The dose rate was 8.47 e^-^/Å^2^/s at the specimen plane. The total exposure time was 6 s with intermediate frames recorded every 0.2 s, giving a total of 30 frames per image. The total accumulated dose was ∼51 e^-^/Å^2^.

### Single-particle cryo EM-image Processing

The frames from each exposure were aligned to compensate for drift and beam-induced motion and summed to a single micrograph using MotionCor2 [46] in Relion 2.1 [47, 48]. Single particles were manually picked using Relion 2.1, and defocus and astigmatism were estimated with CTFFIND4 [49] using movie frames without dose weighting. For the sample containing the inner membrane rings and the export apparatus, 104,021 particles were extracted from 2×2 down sampled, 1,847 motion corrected and dose-weighted micrographs. The extracted box size was 280×280 pixels. The particles were subjected to rounds of reference-free 2D classification in Relion 2.1 with poorly defined classes discarded at each 2D classification stage. Particles selected after the 2D classification (51,833 particles) were further classified in 3D into 6 classes using as a reference a density map calculated from the known structure of the IR1 (accession code 5TCP), which was low pass-filtered to a resolution of 60 Å. At the end of the 3D classification step, a subset of 25,129 double-layered particles was retained for 3D auto-refinement in Relion 2.1. This resulted in a 3D reconstruction map of the double-layered particles with a reported overall resolution of 3.9 Å estimated based on the Fourier shell correlation (FSC) using the 0.143 cut-off criterion. From the 25,129 double-layered particles set, the signal corresponding to IR1 only was subtracted. The IR1 signal subtraction was followed by a new 2D classification round to further screen the resulting 25,129 particle stack representing IR1 images only. At the end of this 2D classification procedure, a subset of 23,433 particles representing IR1 images only was selected and subjected to 3D auto-refinement with a soft mask, resulting in a 3D reconstruction map with a resolution of 3.3 Å as assessed by the FSC. Local resolution variations were estimated from two half data maps using ResMap [50]. For the sample of inner membrane rings without the export apparatus, 43,040 particles were extracted from 1,567 motion corrected and dose-weighed micrographs. The extracted box size was 360×360 pixels. The extracted particles were subjected to several rounds of 2D classification in Relion 2.1 with poorly defined classes discarded at each 2D classification stage. After the 2D classification 27,433 particles were selected and further classified in 3D into 6 classes. At the end of the 3D classification procedure, a set of 13,732 particles representing the most homogeneous subset of 23-membered rings was retained for refinement in Relion 2.1. The resolution of the final map was estimated to be 5.2 Å as assessed by the FSC. This set of particles was subsequently subjected to a new 2D classification round to select for good quality IR1 images. A subset of 10,674 particles representing IR1 only was 3D reconstructed imposing 23-fold symmetry. This improved the resolution of the IR1 to 3.53 Å as assessed by the FSC. Local resolution estimations were calculated from two half data maps using ResMap [50].

### Single particle cryo EM model fitting, refinement and validation

The cryo-EM structure of the IR1 (PDB code: 6DUZ) was fitted by using rigid body fitting into the local resolution filtered 3.3 Å IR1 map, using UCSF Chimera (Pettersen et al., 2004). This was followed by multiple rounds of structure refinement using PHENIX real-space refinement (Adams et al., 2010) to improve the model geometry and fit to the 3.3 Å IR1 map. Geometric and secondary-structure element restraints were maintained throughout the refinement to minimize over-fitting. The outliers were corrected in Coot [51]. A model for the 23-membered ring was constructed in UCSF Chimera (Pettersen et al., 2004) using a monomer subunit from the PDB entry 6DUZ which was sequentially fitted and rigid body refined into the 23-subunits of the local resolution filtered 3.53 Å IR1 map. The resulting 23-membered ring model was further optimized using PHENIX real-space refinement (Adams et al., 2010) with geometric and secondary-structure element restraints maintained throughout the refinement to minimize over-fitting. The outliers were corrected in Coot [51]. Overall model quality and geometry outliers for final models were reported using MolProbity [52]. Details about the maps and final models are presented in S3 Table.

### Electron microscopy and negative staining

Four-microliter drops of a sample containing needle complex preparations were applied to glow-discharged, carbon-coated copper grids and after 30 seconds, samples were blotted with filter paper. Subsequently, 2 µl of a 2% (wt/vol) solution of uranyl acetate was added to the grids and the excess stain was removed by blotting with filter paper. Images of negative-stained samples were recorded with a Gatan 4k-by-4k CCD camera on an FEI Tecnai T12 (120KV) microscope. A total of 71 micrographs with 7,279 selected particles were imaged for samples containing needle complexes from wild type S. Typhimurium, 52 micrographs with 4,346 selected particles for the PrgK^Δ94-95^ mutant, 91 micrographs with 4,943 selected particles for the PrgK^Δ93-95^ mutant, and 192 micrographs with a total of 2,936 selected particles for the PrgK^Δ93-96^ mutant. Particle picking and several cycles of classification and averaging were performed with Relion 2.1 [47, 48].

### Cryo-ET sample preparation

Cryo-ET samples obtained from wild type S. Typhimurium and several export apparatus mutants (S4 Table) were prepared as previously described [25]. Briefly, minicell producing bacterial strains were grown overnight at 37 °C in LB containing 0.3M NaCl. Fresh cultures were prepared from a 1:100 dilution of the overnight culture and then grown at 37 °C to late log phase in the presence of ampicillin (200 µg/mL) and L-arabinose (0.1%) to induce the expression of regulatory protein HilA and thus increase the number of injectisomes partitioning to the minicells. To enrich for minicells, the culture was centrifuged at 1,000 ×g for 5 min to remove normal-size bacterial cells, and the supernatant fraction was further centrifuged at 20,000 ×g for 20 min to collect the minicells. The minicell pellet was resuspended in PBS and mixed with 10 nm colloidal gold particles and deposited onto freshly glow-discharged, holey carbon grids for 1 min. The grids were blotted with filter paper and rapidly frozen in liquid ethane, using a gravity-driven plunger apparatus.

### Cryo-ET data collection and reconstruction

The frozen-hydrated specimens were imaged with a 300kV electron microscope (Titan Krios, Thermo Fisher Scientific) equipped with an energy filter (Gatan) with VPP; the effective pixel size was 2.7 Å. Images were collected automatically using SerialEM [53] in dose fraction mode. The cumulative electron dose for each single-axis tilt series was ∼50e/Å^2^, distributed over 35 images and covering an angular range of −51° to +51° with increments of 3°. Raw images were processed using MotionCor2 [46]. The tilt series were aligned automatically using IMOD [54]. Tomograms were generated by using TOMO3D [55]. In total, 1,266 tomograms (3 × 3,600 × 710 x 1,800 voxels) were generated for detailed examination of the export apparatus in several mutants (Table S4).

### Sub-tomogram analysis

Sub-tomogram analysis was accomplished as described previously to analyze the injectisomes [25]. Briefly, we first visually identified the injectisomes on each minicell. Two coordinates along the needle were used to estimate the initial orientation of each particle assembly. For initial analysis, 4 × 4 × 4 binned sub-tomograms (128 × 128 × 128 voxels) of the intact injectisome were used for alignment and averaging by using the tomographic package I3 [56, 57]. Then multivariate statistical analysis and hierarchical ascendant classification were used to analyze the injectisome [58].

### Three-Dimensional Visualization and Modeling

IMOD [54] was used to take snapshots of 2D slices from 3D tomograms and UCSF Chimera [59] was used for surface rendering of 3D averaged structures. The pseudo atomic model was built by docking the atomic structures of an InvA homolog (PDB-4A5P)[30], SpaP/SpaQ/SpaR homologs (PDB-6F2E), [27], InvC/InvI homologs (PDB-6NJP) [60], and the needle complex (PDB-5TCR) [61]. Animations were generated using UCSF ChimeraX and edited with iMovie.

## Acknowledgements

This work was supported by Grants AI030492 (to J. E. G.), AI126158 (to M. L.-T.), and GM107629 and AI087946 (to J. L.) from the National Institutes of Health.

## Authors’ contribution

C. B. carried out the single particle cryo-EM analyses, W. L. and J. L. carried out the cryo-ET analyses, M. L.-T. prepared all the samples and bacterial strains and carried out the functional analyses, J. L. and J. E. G. designed and directed the studies. J. E. G. wrote the manuscript with contributions from C. B., M. L. T. and J. L.

## Supplementary Materials

**S1 Fig.**
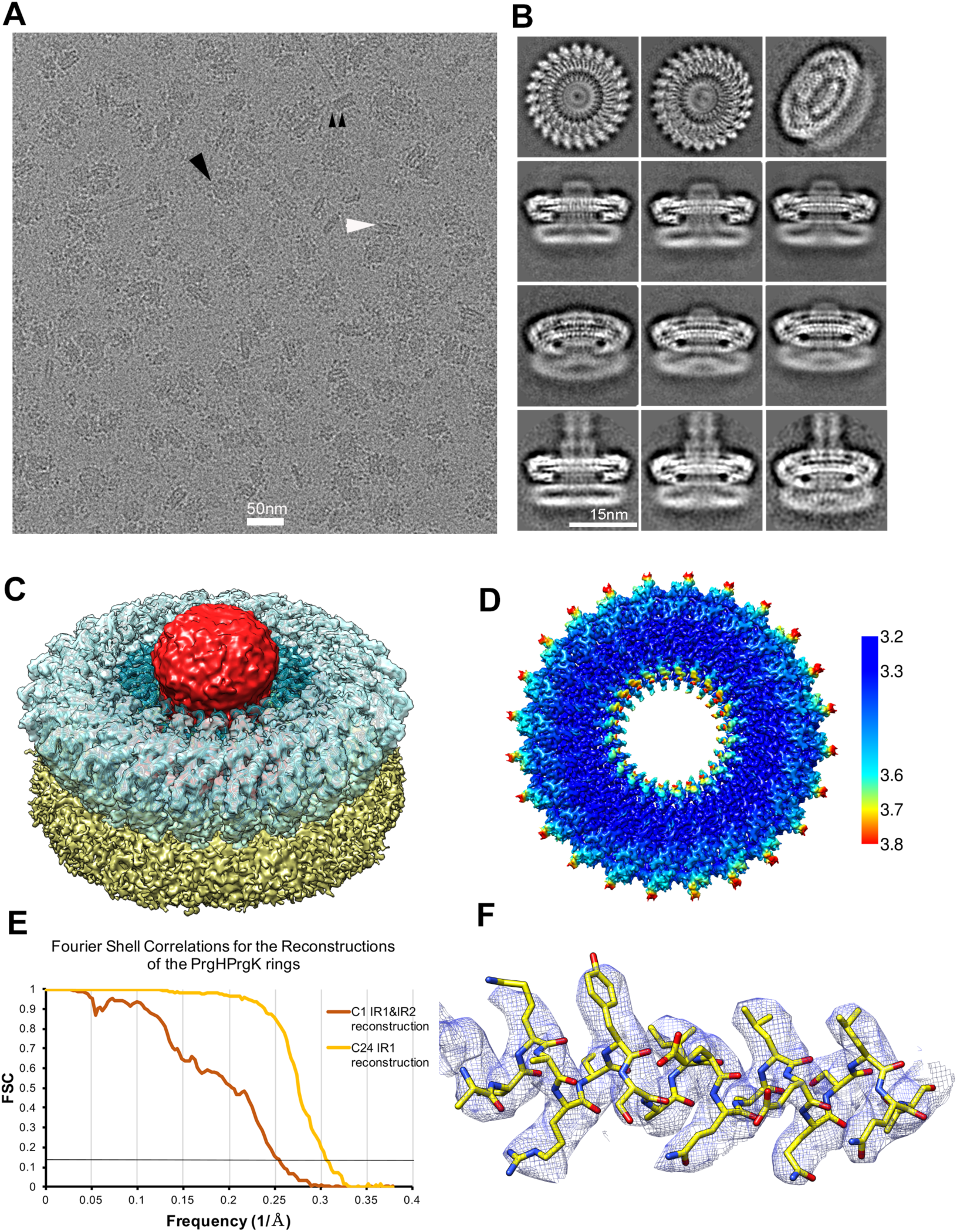
Cryo-EM structure of the needle complex IR1 and IR2 containing the core components of the export apparatus. (A) Representative cryo-EM micrograph. (B) Subset of selected 2D class averages. (C) 3D structure of IR1 (blue), IR2 (yellow) and the core components of the export apparatus (red). (D) Cryo EM structure of the IR1 colored according to the local resolution (in Å). (E) Fourier shell correlations (FSC) of the reconstructions of the IR1 and IR2 rings. (F) Representative cryo-EM local densities with refined atomic models for residues 100-119 of PrgK.

**S2 Fig.**
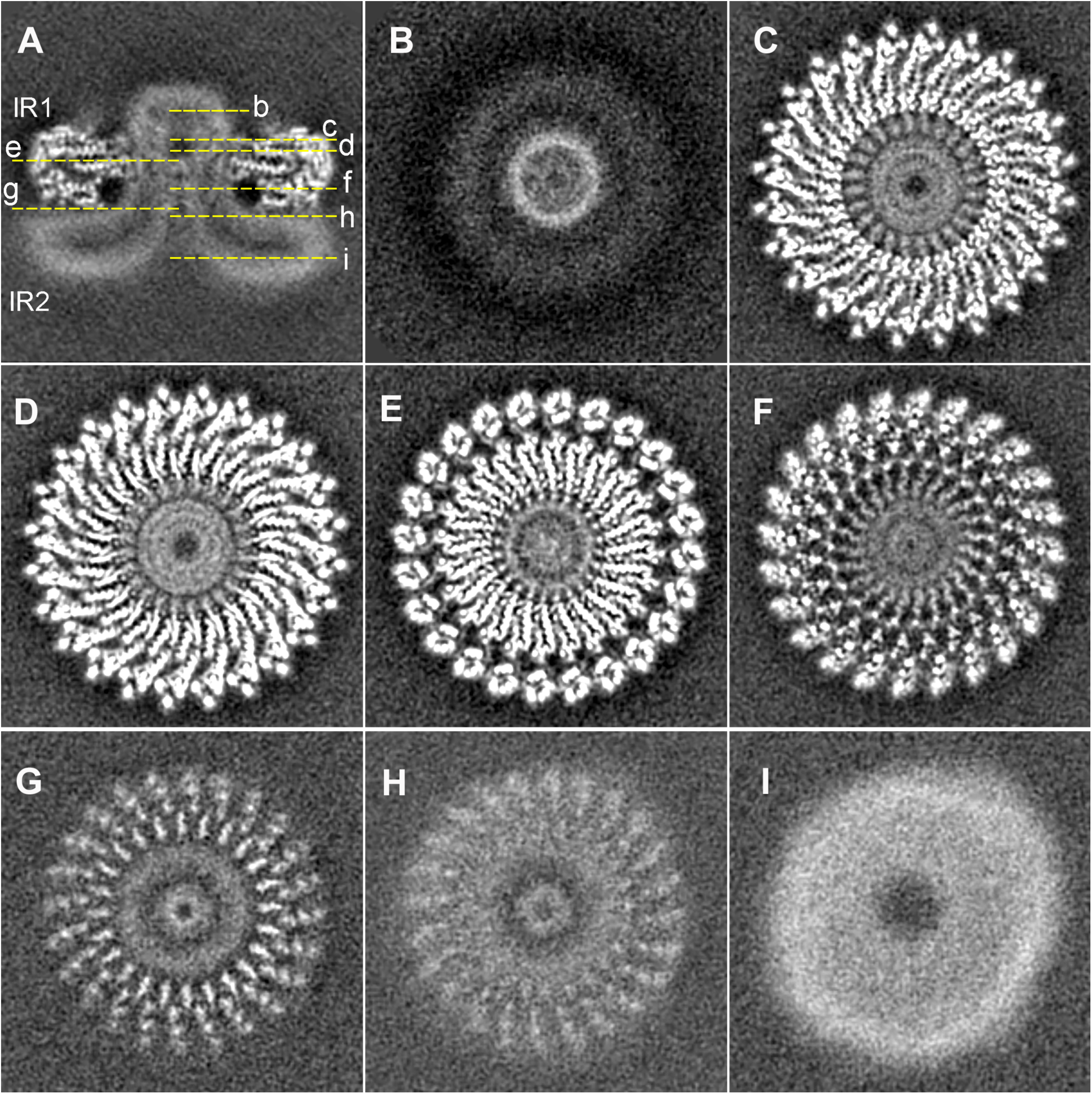
Horizontal sections through the 3D reconstructions of the needle complex IR1 and IR2 containing the core components of the export apparatus. The position of the sections depicted in panels B through I are indicated in panel A.

**S3 Fig.**
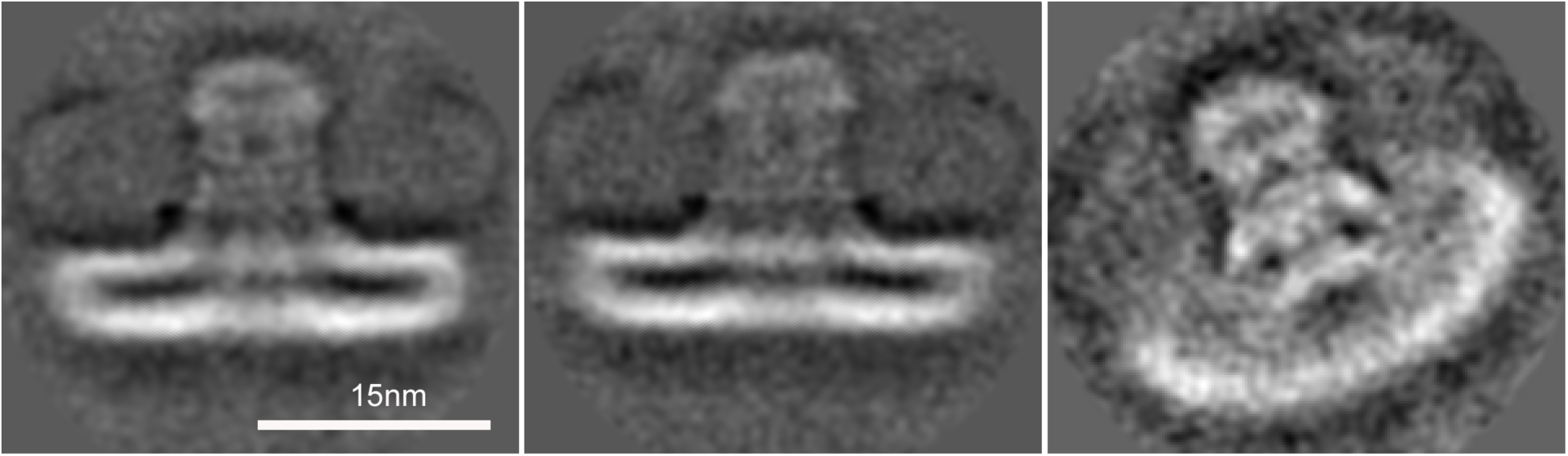
Densities corresponding to IR2, export apparatus, and associated membranes after subtraction of the densities associated with IR1.

**S4 Fig.**
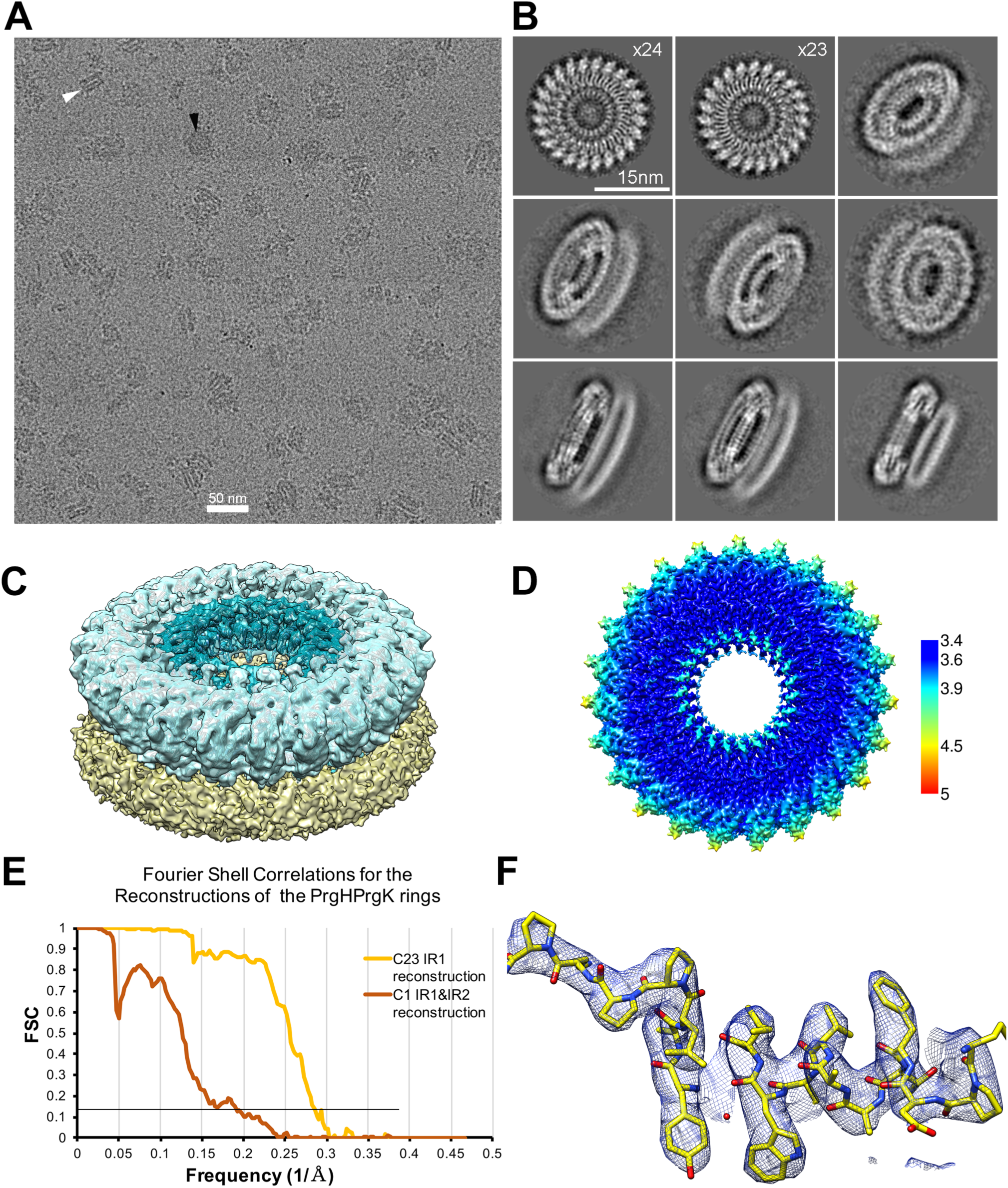
Cryo-EM structure of the needle complex IR1 and IR2 assembled in the absence of the export apparatus showing 23-fold symmetry (A) Representative cryo-EM micrograph. (B) Subset of selected 2D class averages. (C) 3D structure of IR1 (blue), and IR2 (yellow). (D) Cryo EM structure of the IR1 colored according to the local resolution (in Å). (E) Fourier shell correlations (FSC) of the reconstructions of the IR1 and IR2 rings. (F) Representative cryo-EM local densities with refined atomic models for residues 63-81 of PrgK.

**S5 Fig.**
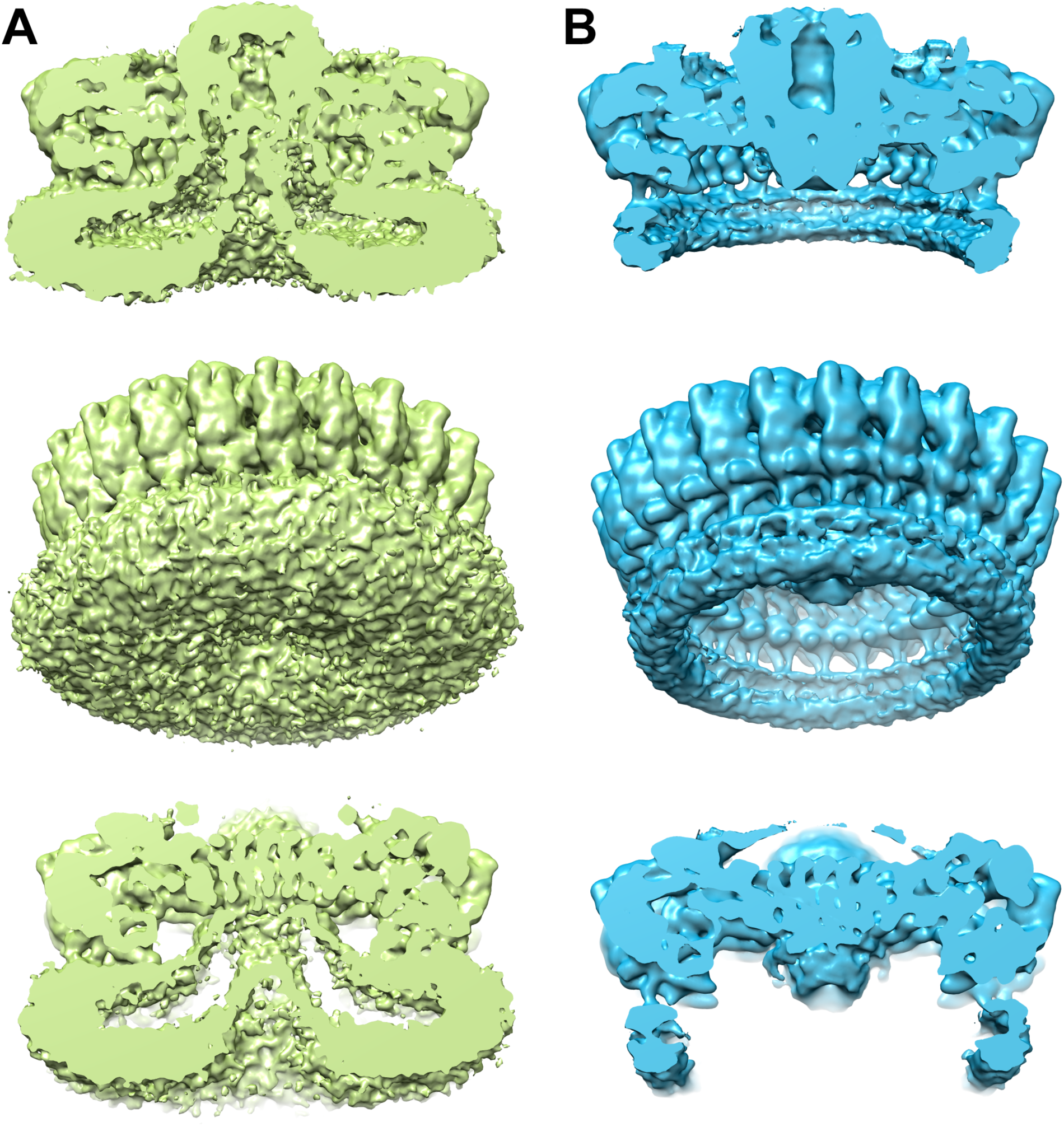
Comparison of the 3D reconstructions of the structure reported in this study (green) (A) with a previously reported structure (accession numbers EMD-8398 and 5TCP) (B) isolated under conditions that remove the associated membranes.

**S6 Fig.**
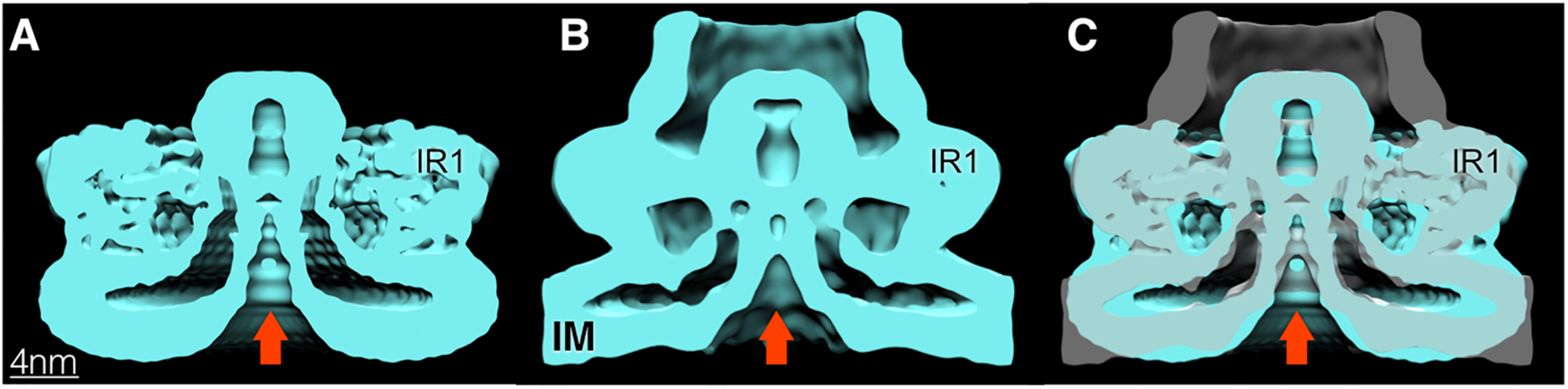
Comparison of the cryo-EM structure of the isolated needle complex IR1 and IR2 containing the core components of the export apparatus (A) with the cryo ET structures of the type III secretion needle complex from a *S*. Typhimurium *ΔinvA* mutant strain (B). The overlaid structures are shown in (C).

**S7 Fig.**
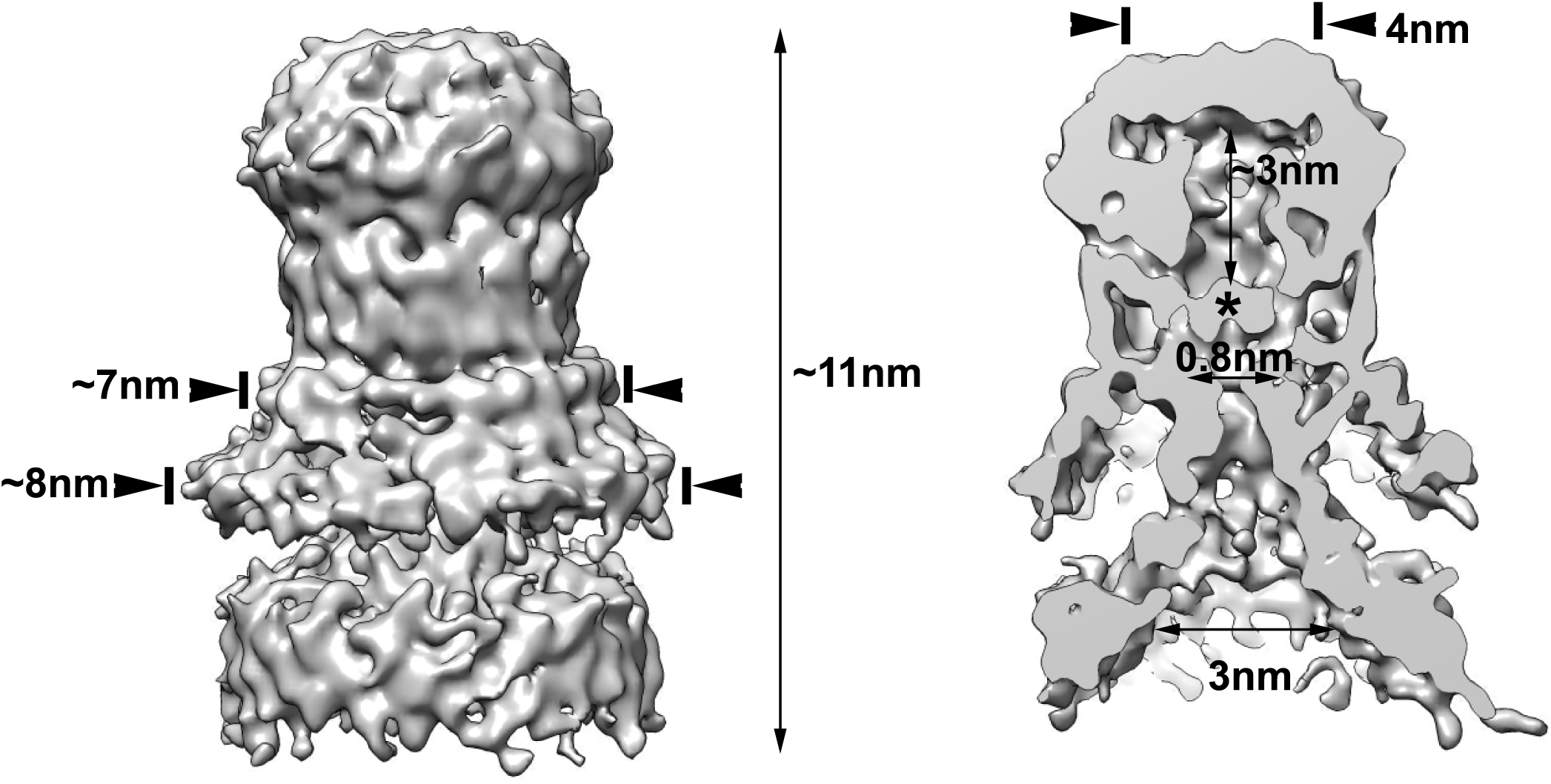
3D reconstructions of the core components of the export apparatus and associated membranes after masking the densities corresponding to the IR1 and part of the IR2. In a cut-away view through the 3D reconstruction, an asterix denotes a density that separates the central cavity from the cytoplasmic side of the structure. Relevant dimension are indicated.

**S8 Fig.**
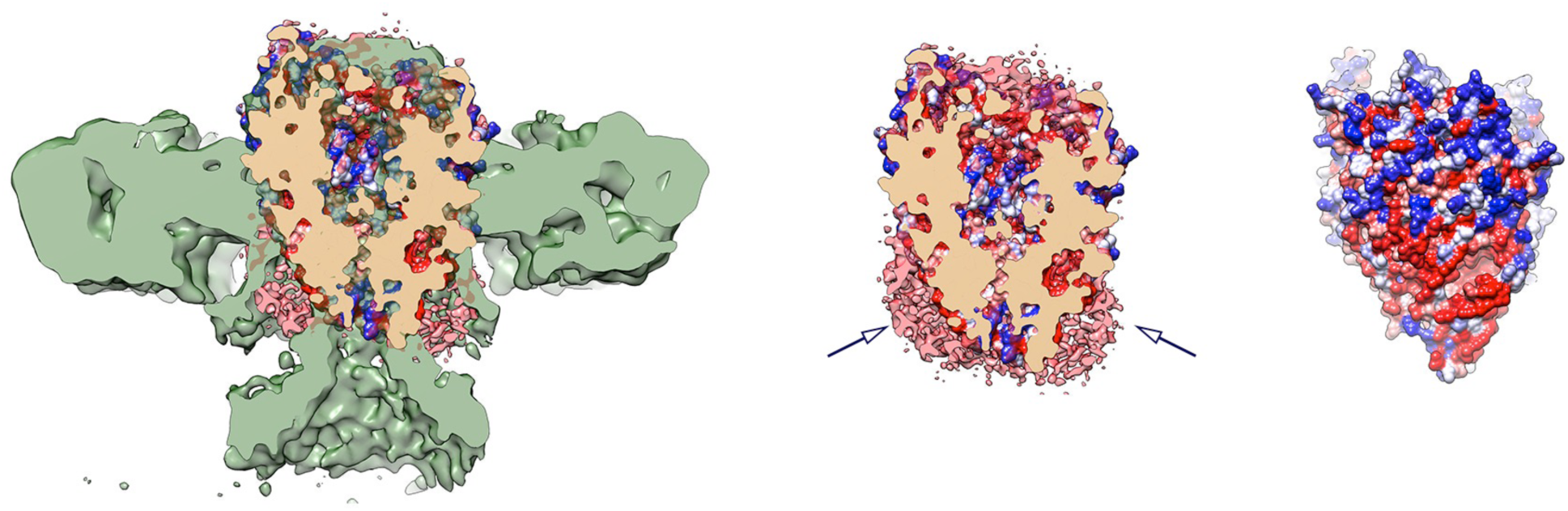
Overlay of the cryo-EM structure of the needle complex IR1 and IR2 containing the core components of the export apparatus (green) with the structure of the isolated FliP/FliQ/FliR complex (PDB 6F2E). The surface of the FliP/FliQ/FliR complex surface is colored according to hydrophobicity (red -hydrophobic; blue-charged). The location of the detergent belt (in pink, central image) is indicated by arrows.

**S9 Fig.**
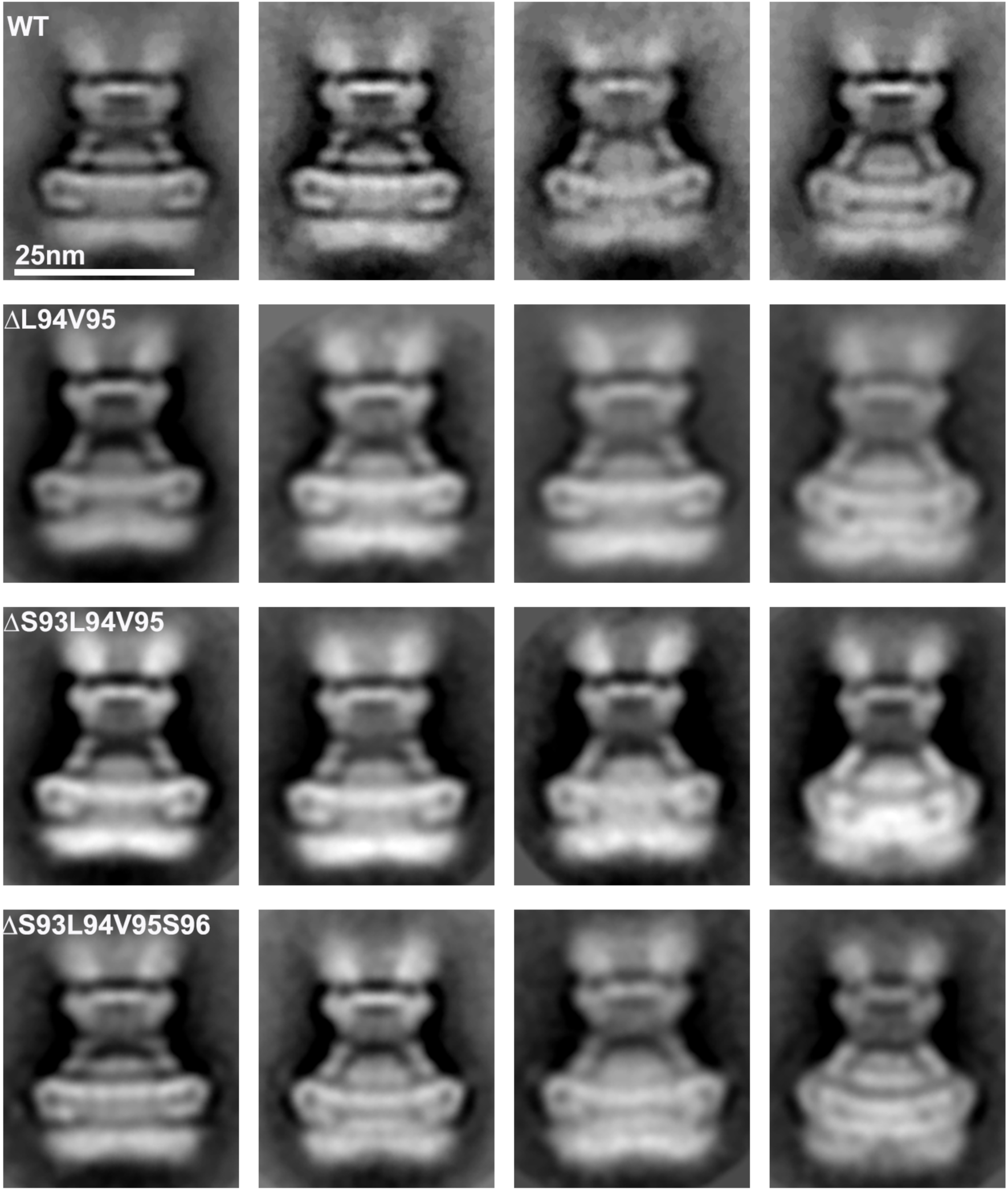
Mutations in a periplasmic domain of the needle complex protein PrgK that interacts with the core components of the export apparatus do not affect needle complex base structure or export apparatus deployment. Shown are representative class averages of needle complexes isolated from *S*. Typhimurium expressing wild type PrgK or the indicated PrgK deletion mutants.

**S10 Fig.**
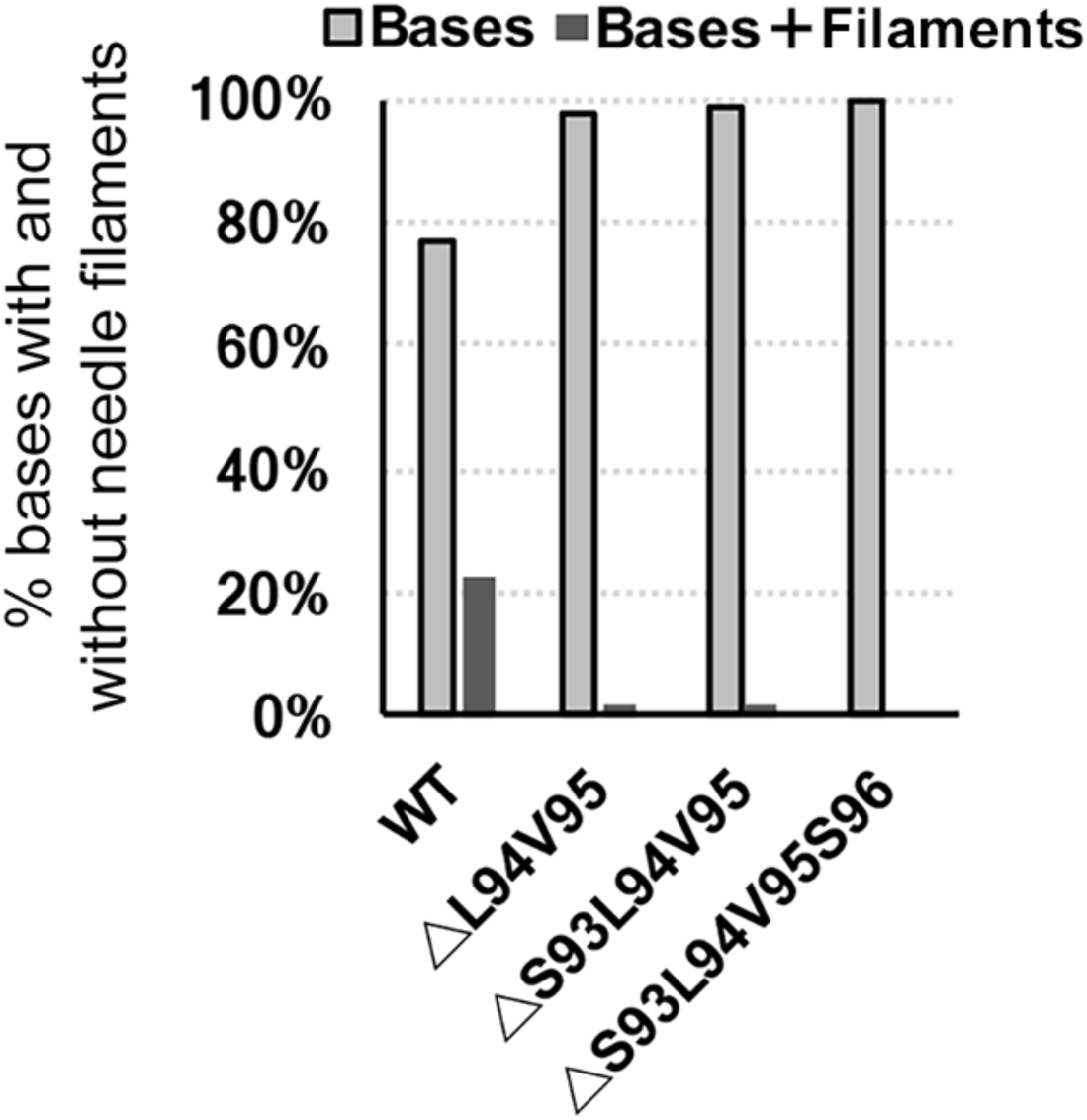
Deletions mutations in a periplasmic domain of the needle complex protein PrgK impairs needle filament assembly. The graph shows the percentage of needle complex bases with (bases) or without (bases + filaments) needle filaments.

**S11 Fig.**
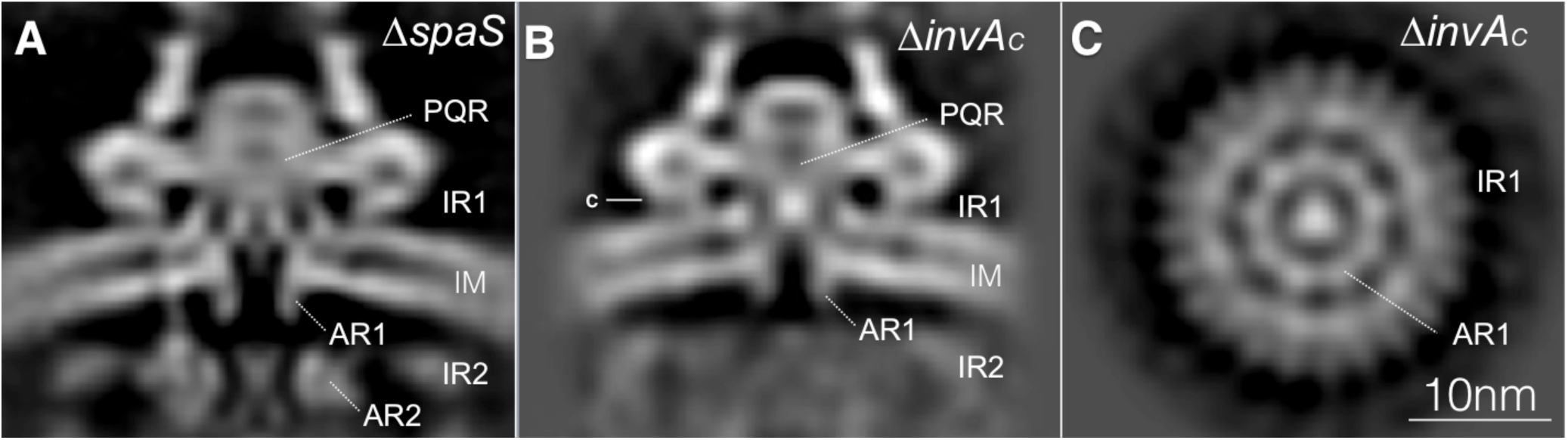
Vertical sections of the cryo ET *in situ* structures of injectisomes obtained from S. Typhimurium *ΔspaS* (A), or from a strain expressing a mutant form of InvA that lacks its last 328 amino acids (B). Also shown is a horizontal section of the latter structure (C).

**S12 Fig.**
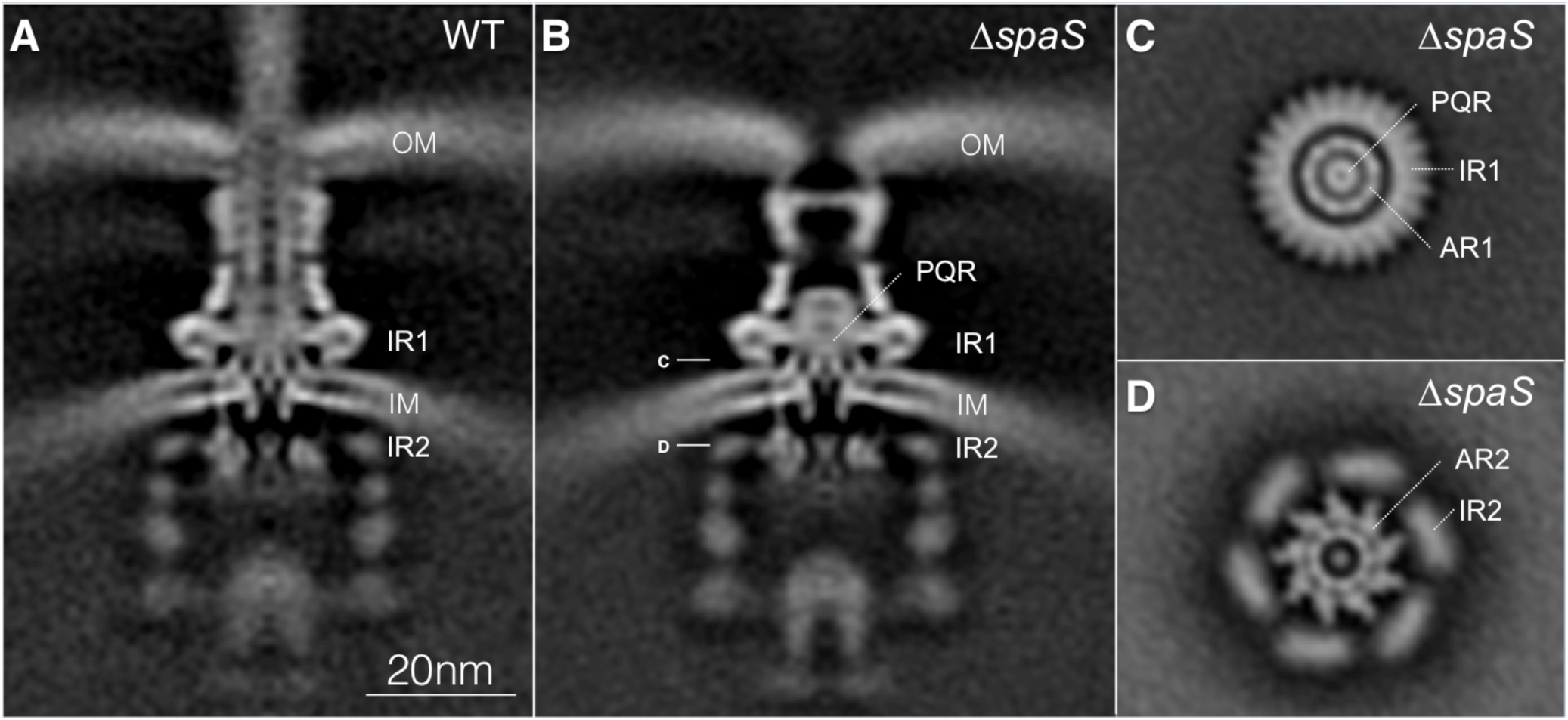
Vertical sections of the cryo ET *in situ* structures of injectisomes obtained from wild type S. Typhimurium (A) or ΔspaS mutant strain (B), Also shown are a horizontal sections of the latter structure strain through the planes indicated in panel B (C and D). OM: bacterial outer membrane; IM: bacterial inner membrane; IR1 and IR2: inner rings 1 and 2 of the T3SS needle complex; PQR: SpaP/SpaQ/SpaR complex; AR1 and AR2: membrane proximal (AR1) and distal (AR2) cytoplasmic rings of InvA.

**Table S1:**
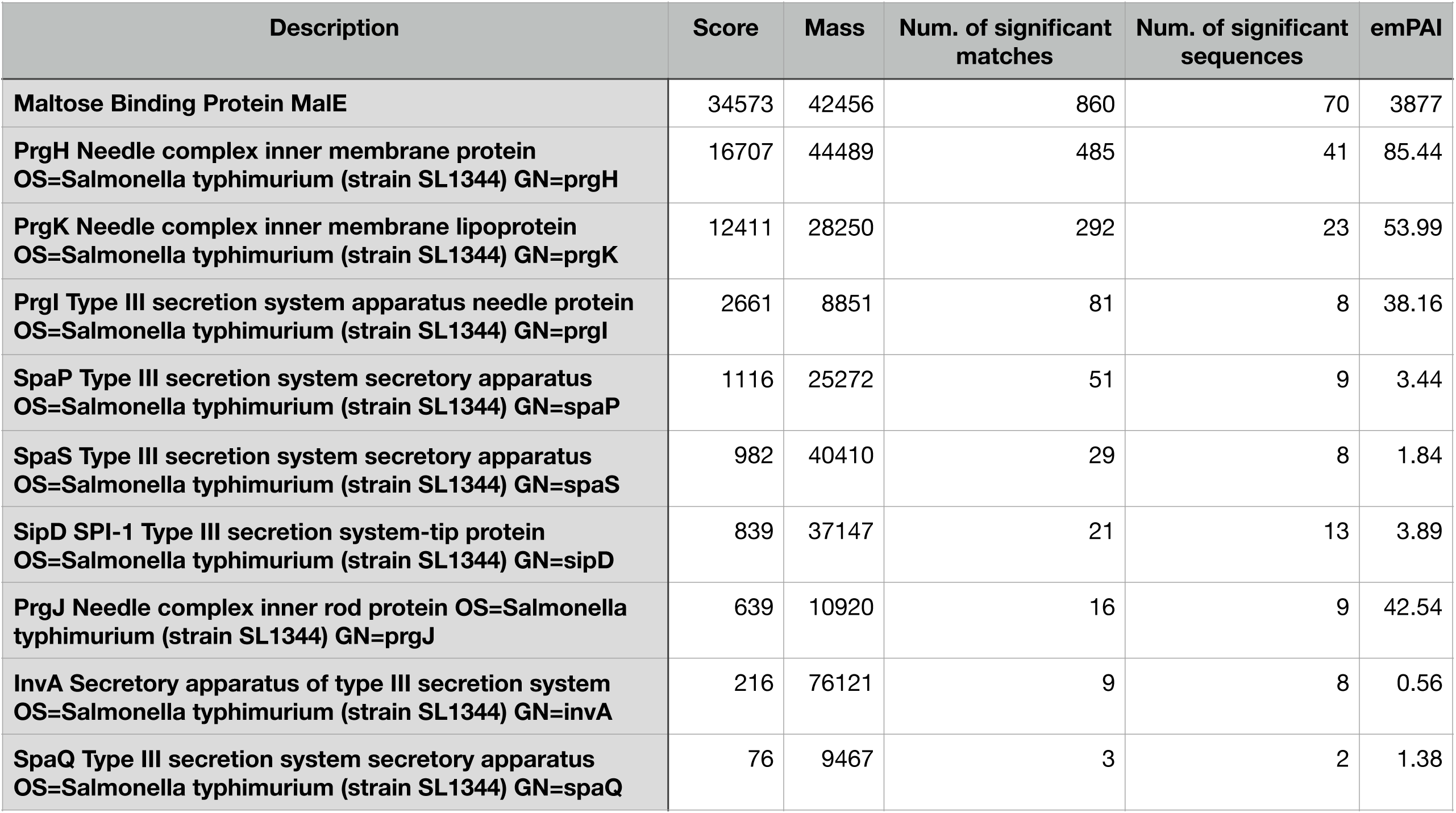
LC-MS/MS analysis of needle complex preparation

| Description | Score | Mass | Num. of significant matches | Num. of significant sequences | emPAI |
| --- | --- | --- | --- | --- | --- |
| **Maltose Binding Protein MalE** | 34573 | 42456 | 860 | 70 | 3877 |
| **PrgH Needle complex inner membrane protein**<br>**OS=Salmonella typhimurium (strain SL1344) GN=prgH** | 16707 | 44489 | 485 | 41 | 85.44 |
| **PrgK Needle complex inner membrane lipoprotein**<br>**OS=Salmonella typhimurium (strain SL1344) GN=prgK** | 12411 | 28250 | 292 | 23 | 53.99 |
| **PrgI Type III secretion system apparatus needle protein**<br>**OS=Salmonella typhimurium (strain SL1344) GN=prgI** | 2661 | 8851 | 81 | 8 | 38.16 |
| **SpaP Type III secretion system secretory apparatus**<br>**OS=Salmonella typhimurium (strain SL1344) GN=spaP** | 1116 | 25272 | 51 | 9 | 3.44 |
| **SpaS Type III secretion system secretory apparatus**<br>**OS=Salmonella typhimurium (strain SL1344) GN=spaS** | 982 | 40410 | 29 | 8 | 1.84 |
| **SipD SPI-1 Type III secretion system-tip protein**<br>**OS=Salmonella typhimurium (strain SL1344) GN=sipD** | 839 | 37147 | 21 | 13 | 3.89 |
| **PrgJ Needle complex inner rod protein OS=Salmonella typhimurium (strain SL1344) GN=prgJ** | 639 | 10920 | 16 | 9 | 42.54 |
| **InvA Secretory apparatus of type III secretion system**<br>**OS=Salmonella typhimurium (strain SL1344) GN=invA** | 216 | 76121 | 9 | 8 | 0.56 |
| **SpaQ Type III secretion system secretory apparatus**<br>**OS=Salmonella typhimurium (strain SL1344) GN=spaQ** | 76 | 9467 | 3 | 2 | 1.38 |

**Table S2:** Strains used in this study

| Strain | Genotype | Reference (PMID) |
| --- | --- | --- |
| SB3083 | *MBP-prgH ΔinvG flhD::Tn10* | This study |
| SB3098 | *MBP-prgH ΔinvG ΔspaPQRS flhD::Tn10* | This study |
| SB3079 | *MBP-prgH flhD::Tn10* | This study |
| SB3581 | *MBP-prgH prgKΔL94V95 flhD::Tn10* | This study |
| SB3582 | *MBP-prgH prgKΔS93L94V95 flhD::Tn10* | This study |
| SB3583 | *MBP-prgH prgKΔS93L94V95S96 flhD::Tn10* | This study |
| SB1780 | *minD::cat* | 23481398 |
| SB3112 | *ΔinvA minD::cat* | 28283062 |
| SB3113 | *3xFspaO ΔspaPQRS minD::cat* | 28283062 |
| SB3565 | *ΔspaS minD::cat* | This study |
| SB3592 | *invAΔ357-685aa-3xF minD::cat* | This study |

**Table S3.**
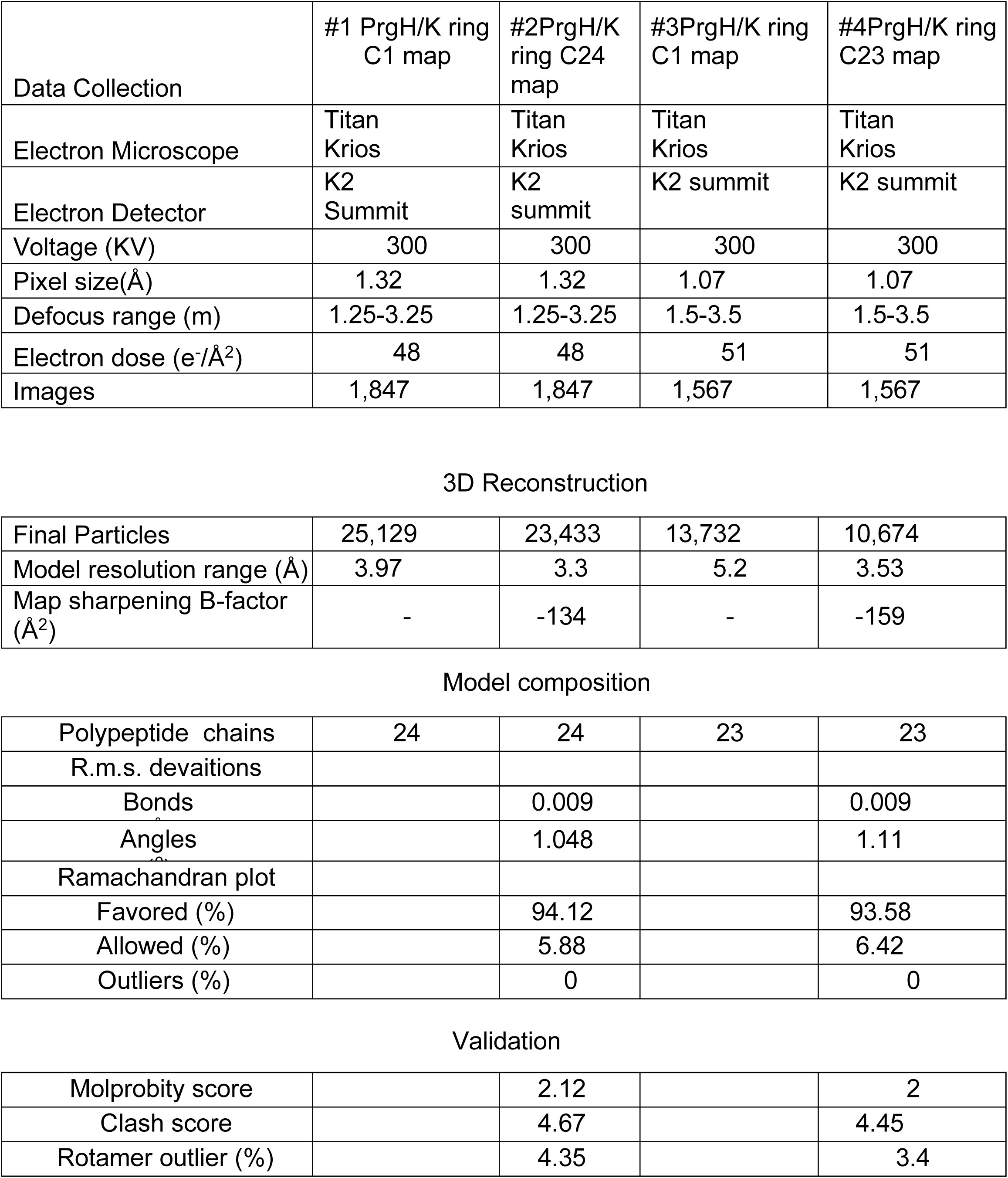
Cryo EM data

|  | #1 PrgH/K ring<br>C1 map | #2PrgH/K<br>ring C24<br>map | #3PrgH/K ring<br>C1 map | #4PrgH/K ring<br>C23 map |
| --- | --- | --- | --- | --- |
| Data Collection |  |  |  |  |
| Electron Microscope | Titan<br>Krios | Titan<br>Krios | Titan<br>Krios | Titan<br>Krios |
| Electron Detector | K2<br>Summit | K2<br>summit | K2 summit | K2 summit |
| Voltage (KV) | 300 | 300 | 300 | 300 |
| Pixel size(Å) | 1.32 | 1.32 | 1.07 | 1.07 |
| Defocus range (m) | 1.25-3.25 | 1.25-3.25 | 1.5-3.5 | 1.5-3.5 |
| Electron dose (e<sup>-</sup>/Å<sup>2</sup>) | 48 | 48 | 51 | 51 |
| Images | 1,847 | 1,847 | 1,567 | 1,567 |
| 3D Reconstruction |  |  |  |  |
| Final Particles | 25,129 | 23,433 | 13,732 | 10,674 |
| Model resolution range (Å) | 3.97 | 3.3 | 5.2 | 3.53 |
| Map sharpening B-factor<br>(Å<sup>2</sup>) | - | -134 | - | -159 |
| Model composition |  |  |  |  |
| Polypeptide chains | 24 | 24 | 23 | 23 |
| R.m.s. deviations |  |  |  |  |
| Bonds |  | 0.009 |  | 0.009 |
| Angles |  | 1.048 |  | 1.11 |
| Ramachandran plot |  |  |  |  |
| Favored (%) |  | 94.12 |  | 93.58 |
| Allowed (%) |  | 5.88 |  | 6.42 |
| Outliers (%) |  | 0 |  | 0 |
| Validation |  |  |  |  |
| Molprobity score |  | 2.12 |  | 2 |
| Clash score |  | 4.67 |  | 4.45 |
| Rotamer outlier (%) |  | 4.35 |  | 3.4 |

**Table S4:** Number of tomograms of minicells and sub-tomograms of injectisomes processed in this study

| Strain | Relevant Genotype | Tomograms | Sub-tomograms |
| --- | --- | --- | --- |
| SB1780 | *wild-type* | 326 | 6523 |
| SB3112 | $\Delta invA$ | 275 | 4165 |
| SB3113 | $\Delta spaPQRS$ | 241 | 2368 |
| SB3565 | $\Delta spaS$ | 316 | 5882 |
| SB3592 | *invA*Δ | 108 | 992 |
| Total: |  | 1266 | 19,930 |

## Video captions

**S1 Video**. Docking of the structure of the core components of the export apparatus FliP/FliQ/FliR (surface rendered according to hydrophobicity (red, hydrophobic; blue, charge) onto the cryo EM structure of isolated needle rings of the SPI-1 S. Typhimurium T3SS injectisome containing the export apparatus.

**S2 Video.** Localization of the export apparatus component InvA (depicted in pink) relative to the core components of the export apparatus (depicted in yollow) and other structures of the type III secretion injectisome.

**S3 Video.** Depiction of the conduit delimited by the InvA cytoplasmic rings that leads to the entrance of the gate of the export apparatus.

